# A water-pinning hotspot drives templated tau aggregation

**DOI:** 10.64898/2026.08.04.742883

**Authors:** Chung-Ta Han, Karen Tsay, Samuel Lobo, Saeed Najafi, Kieran Monahan, Erica Hong, M. Scott Shell, Joan-Emma Shea, Songi Han

## Abstract

Tau assembles into fibrillar aggregates that are pathological hallmarks of a group of neurodegenerative diseases collectively called tauopathies. Templated aggregation of naïve tau to seeding-competent fibrils that proceed from cell to cell is a key driver of prion-like progression of tauopathies. This study tests the hypothesis that tau, an intrinsically disordered protein (IDP), achieves in-register stacking to form seed-competent fibrils by a pinning action of tau to each other and/or the seed surface via a single dominant hotspot to avoid mismatch in tau stacking to fibrils. Structured solvation water has been proposed to be a signature of such hotspots at both the tau fibril-end surface and soluble tau monomers. Although jR2R3-P301L tau exhibits a heterogeneous hydration landscape in its intrinsically disordered state, with enhanced water structuring near the P301L mutation site, it is unclear whether a localized hotspot exists at the fibril end surface and surface water facilitates the initial contacts in templated aggregation. Using rapid ^1^H-^15^N SOFAST-HMQC NMR to track seed-induced aggregation of jR2R3-P301L in real time, complemented by molecular dynamics simulation of fibril surface hydration, we identify a residue-specific pinning hotspot that is prone to dewetting followed by sequential folding and incorporation of the remaining segment in a two-step dock-and-lock process. Site-specific spin labeling further demonstrates that blocking this pinning hotspot disrupts templated aggregation, leading to shorter fibrils. The identification of a dominant pinning site will facilitate the rational design of binders to effectively disrupt fibril extension or serve as diagnostic or therapeutic strategies.

**Significance Statement:** Tau proteins must align and stack precisely with existing fibril ends to propagate pathology, yet the molecular signature that initiates and ensures this in-register alignment has been unclear. This study shows recruitment begins at a single, structurally well-defined contact site on the fibril surface that is enriched in release-prone hydration water. These findings reveal that water-release-prone hotspots, rather than the well-known amyloid-core regions alone, govern the initial steps in templating seeding and can hence be blocked, opening a new path for identifying therapeutic targets that could slow the progression of tau-related diseases.

## Introduction

Tau is an intrinsically disordered protein (IDP) that normally remains associated with microtubules to support their stability, but can misfold and aggregate into insoluble amyloid fibrils under neurodegenerative disease conditions collectively known as tauopathies (1). The accumulation of tau aggregates, particularly in the form of neurofibrillary tangles (NFTs) is clinically diagnostic of tauopathies and is consistently observed as a defining hallmark of tauopathies according to postmortem pathologies. Recent advances in cryogenic electron microscopy (cryo-EM) have resolved the atomic structures of tau fibrils across many of these tauopathies, including Alzheimer’s disease (AD) (2, 3), Pick’s disease (PiD) (4), chronic traumatic encephalopathy (CTE) (5), corticobasal degeneration (CBD) (6), progressive supranuclear palsy (PSP) (7), argyrophilic grain disease (AGD) (8), aging-related tau astrogliopathy (ARTAG) (9), globular glial tauopathy (GGT) (9), and frontotemporal dementia and parkinsonism linked to chromosome 17 (FTDP-17) (10), revealing that tau filaments form disease-specific tertiary and quaternary protein folds. These fibril structures are diagnostic hallmarks of tauopathies, but currently mainly solved for tau fibrils derived from postmortem tissues. These would be valuable diagnostic targets if key early events of their disease-specific folding, assembly and propagation are identified.

The central hypothesis of this study is that the earliest assembly of tau and the subsequent growth of seeding-competent fibrils are driven by a single, highly localized interaction hotspot. Such a site would provide the initial alignment needed for tau, an IDP, to adopt the in-register β-sheet stacking required for fibril formation. A dominant hotspot of this kind would also represent an attractive target for blocking the prion-like propagation of tau aggregation. We propose that this principle applies broadly to tau constructs capable of forming seeding-competent fibrils, provided that reliable in-register stacking along the fibril axis can be achieved. To test this idea, we set out to experimentally capture the residue-level signature of such a hotspot using a model tau peptide previously shown to adopt a disease-specific fold and assemble into highly regular, seed-competent fibrils. The tau peptide of choice includes a hexapeptide segment ^306^VQIVYK^311^ located in the R3 domain (PHF6) of the microtubule-binding region of tau, a sequence that forms part of the core in all resolved tau fibril structure. The construct is extended to include the short segment that folds back to pair with PHF6 in the conserved strand-loop-strand (SLS) motif shared between 4R tauopathies (CBD, PSP, GGT, AGD, LNT, and FTDP-17), connected through a short loop in a U-shaped fold (9). Recent studies have demonstrated that short tau fragments containing this U-shaped motif assemble into uniform fibrils that closely recapitulate folds observed in 4R tauopathy fibril structure (11–14). Critically, these minimal fibrils exhibit prion-like properties, serving as seeds that recruit soluble tau and trigger its aggregation, including full-length 0N4R tau (11, 12). However, not all tau peptides containing PHF6 segment form potent seeds, implying that PHF6 alone does not determine seeding efficiency (12, 15). Because seed-induced aggregation underlies the prion-like progression of tauopathies (16), identifying hotspots that enhance this process is essential for developing strategies to effectively block tau fibril propagation (17).

In seed-induced aggregation, pre-existing fibril ends and surfaces lower the energy barrier for tau fibril formation relative to the much slower seed-independent nucleation pathway. The elongation step, often referred to as template seeding, occurs when soluble tau binds to an active fibril end and adopts the same β-sheet structure. In the nucleation pathway, mutations including P301L (12, 18), ΔK280 (19), V337M (20), and R406W (21, 22) accelerate tau self-aggregation. Although the underlying mechanisms remain unclear, prior works on intrinsically disordered tau suggest that these mutations may promote fibril formation by shifting the conformational ensemble toward aggregation-prone states, destabilizing aggregation-incompetent conformers, and/or altering the hydration landscape around the mutated site (12, 23). However, it is unclear whether such mutations also alter the surface properties of the resulting fibril in ways that generate or enhance highly localized hotspots capable of enhancing templated seeding.

A dock–lock model is widely used to describe the templated seeding process, where the docking step involves reversible binding of soluble tau to active fibril ends, and the locking step involves irreversible conformational rearrangements that result in β-sheet formation (24–26). While this model captures key features of fibril elongation, its application to tau templated seeding leaves open a critical question. How do highly disordered IDP tau, with its astronomically large conformational ensembles, consistently find the correct docking site and lock into the rigid cross-β structure? It is energetically impossible for soluble tau to undergo simultaneous conformational alignment and incorporation into the rigid cross-β structure of fibrils in a pre-folded form (27). A more plausible model is that initial contact occurs at a specific preferred hotspot on the fibril end, which dock–lock first, followed by sequential docking, folding, and incorporation of the remaining residues. *The event and signature that this study is after is to identify the existence of such a dominant hotspot serving as the preferred docking site on the tau fibril-end surface and soluble tau under conditions of productive fibril growth in seed-induced aggregation*.

Given that interfacial dewetting governs the formation of intermolecular hydrogen bonds essential for β-sheet stacking, the hydration water at the interface is expected to play an important role in templated seeding (28, 29). Prior simulation studies have shown that structured water with lower entropy on protein surfaces is thermodynamically prone to dewetting, thereby facilitating molecular docking events (30). Consequently, the hydration landscape on the tau monomer and fibril surface should reveal the existence of a dominant hotspot that may drive the initial contact in the templated seeding. Although in a previous study, a heterogeneous hydration landscape has been experimentally observed with a site hydrated with low entropy water identified around soluble tau peptides in their IDP state (12), this was only a necessary condition for the posed hypothesis. Whether the highly localized hotspot on the tau monomer in its IDP state is preserved in the fibril state of this peptide, and whether it is practically the dominant interaction hotspot that first engages in inter-tau interaction, are unknown.

This study aims to directly track, with residue-level resolution, the time course of soluble jR2R3-P301L tau peptide association with preformed tau jR2R3-P301L fibril seeds upon mixing. The association sequence of soluble tau to its fibril surface was tracked by solution state ^1^H-^15^N band-selective optimized flip-angle short-transient heteronuclear multiple quantum coherence (SOFAST-HMQC) NMR to interrogate the kinetics of aggregation with residue-level resolution (31–33). The potential of on-pathway oligomer formation was examined by 2D diffusion-ordered spectroscopy (DOSY) to provide further insight into the molecular-assembly process before the docking of soluble tau jR2R3-P301L peptides to the fibril surface. The hydration dynamics and water structure on the jR2R3-P301L fibril surface were quantified by fully atomistic molecular dynamics (MD) simulations and Overhauser Dynamic Nuclear Polarization (ODNP) relaxometry (34). The goal is to establish the connection between the specific residue(s) on the fibril surface with highly localized structure water and the initial contact event leading to docking followed by locking events in templated seeding. We demonstrate that blocking the identified interaction hotspot on the fibril surface with a spin label attenuates template seeding, but not when spin labels are placed at other sites. This was demonstrated using continuous wave electron paramagnetic resonance (cw-EPR) lineshape analysis, showing that these sites have the potential to serve as strategic targets for therapeutic interventions aimed at stopping the propagation of tau fibrils.

## Results

### Sample Design

This study aims to identify the interaction hotspots that facilitate the templated seeding of tau protein. To achieve this goal, we selected a minimal tau construct that readily forms homogeneous fibrils with a disease-like fold and shows seed competency for prion-like amplification. A 19-amino-acid jR2R3-P301L (^295^DNIKHVPGGGSVQIVYKPV^313^) truncated tau peptide (Fig. 1 A), spanning the R2/R3 domains of full-length tau with a MAPT mutation at site 301 was shown to form seed-competent fibrils that can template and propagate misfolded tau, including full-length 0N4R tau, in a prion-like manner, both in vitro and in cellular models (11, 12). According to the cryo-EM structure, the cross-section of this jR2R3-P301L fibril contains two pairs of protofilaments, with the outer two chains adopting a strand-loop-strand (SLS) motif typical for 4R tauopathies, as found in CBD, LNT, PSP, or GGT, and the inner two chains serving as the counter strands to stabilize the protofilaments (Fig. 1 B) (9). The hypothesis to be tested is whether templated seeding is initiated at a specific hotspot (Fig. 1 C), with the free energy of association provided by the release of localized, tetrahedrally structured and dynamic, vapor-like hydration water (35).

**Fig. 1.**
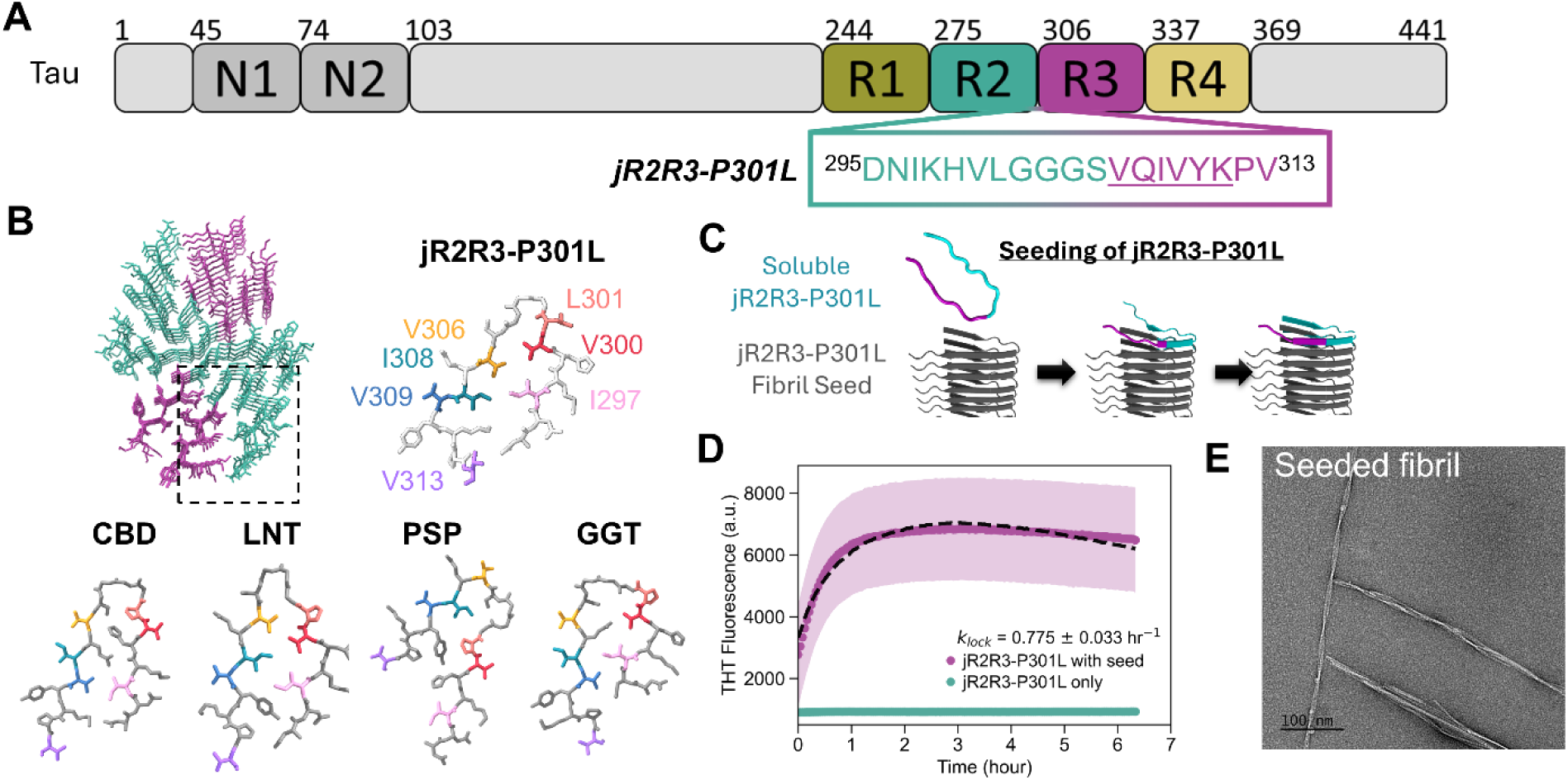
Templated seeding of tau jR2R3-P301L construct: (A) (Upper) The longest isomer of tau protein with four microtubule binding domains colored. (Lower) The sequence of the 19-amino-acid jR2R3-P301L tau peptide and its location in the full-length tau. (B) Fibril structure of jR2R3-P301L (PDB ID #8V1N). The outer chain adopts a strand-loop-strand motif that is shared between 4R tauopathies, including CBD type II (PDB#: 6TJX), LNT (PDB#: 7P6A), PSP (PDB# 7P65), and GGT type I (PDB#: 7P66). The colored residues I297 (pink), V300 (red), L301 (orange), V306 (yellow), I308 (aqua), V309 (blue), and V313 (purple) are the ones enriched with ^13^C and ^15^N for our NMR studies. (C) Schematic of jR2R3-P301L association process based on our hypothesis: with a specific site(s) on the soluble jR2R3-P301L peptide pinned to existing fibril seeds, followed by sequential residue association. (D) ThT fluorescence of jR2R3-P301L templated seeding (soluble peptide plus seed at a 1:1 monomer-equivalent ratio, purple) and the peptide-only control (turquoise). The seeded trace is well described by a single-exponential fit (black), the lock-limited form of the dock-and-lock scheme (Esler et al., 2000), giving k_lock ≈ 0.8 hr⁻¹; under these efficient high-seed conditions the ThT rise reports the locking (cross-β) rate. (E) nsTEM image of the jR2R3-P301L fibril after the templated seeding.

### Seed-induced aggregation of jR2R3-P301L tau

The seed-induced aggregation of jR2R3-P301L tau was first examined by a Thioflavin T (ThT) fluorescence assays and negative stain TEM (Fig. 1 D and E) after mixing soluble ^13^C/^15^N isotope-enriched jR2R3-VLI-P301L peptides with an amount of prepared jR2R3-P301L fibril seeds prepared from an equivalent quantity of peptide (fibril seed preparation is described in Materials and Methods, with TEM images along the preparation workflow shown in Figure S1).

An increase in ThT fluorescence intensity was observed in the sample after initiating templated seeding relative to the seed alone, with an onset of reaction observed immediately after mixing seeds and soluble tau, without a lag phase, and reaching a plateau after 2 hrs (purple, Fig. 1 D), reflecting the low barrier for the seed-induced aggregation. The seeded ThT trace is well described by a single time constant (k_lock_ ≈ 0.8 hr⁻¹, Fig. 1D), which under these efficient high-seed conditions reports the locking rate. In contrast, the control sample containing only jR2R3-P301L tau peptide showed no ThT signal increase (turquoise, Fig. 1 D), suggesting that no self-aggregation of the seed by primary nucleation is occurring over the measured time frame. Negative stain TEM also showed an extension of fibril length after templated seeding (Fig. 1 E, Fig. S1 D) compared to the freshly prepared seeds (Fig. S1 C). No fibril formation was found in the control sample with peptide only (data not shown). This confirms that fibrillization occurred predominantly via templated seeding with negligible self-aggregation, rendering it a well-controlled system to track the molecular detail of templated seeding by NMR, as described next.

### Interaction hotspots in templated seeding of jR2R3-P301L tau identified by NMR

Next, we examine whether soluble tau engages the fibril surface and becomes immobilized in a residue-specific sequence or whether all residues become immobilized simultaneously. We performed a time-resolved series of ^1^H-^15^N SOFAST-HMQC experiments tracking the residue-specific binding sequence in jR2R3-P301L templated seeding by mixing fibrillar seeds made of jR2R3-P301L at natural abundance with jR2R3-VLI-P301L peptides that are ^13^C/^15^N isotope-enriched for seven selected residues of V, L, and I (I298, V300, L301, V306, I308, V309, and V313) spanning the peptide. A 2D ^1^H-^15^N SOFAST-HMQC contour plot of the isotope-enriched peptides in their soluble form resolves seven different ^1^H-^15^N frequency combinations, also referred to as correlations (t = 0 hr, Fig. 2 A). These correlations are observable only when the corresponding residues remain mobile, that is, before their incorporation into fibrils. By acquiring a series of 2D spectrum and tracking the decay of residue-specific peak volumes in ∼15-min intervals after initiating templated seeding, we monitored immobilization of the soluble peptide in real time. If templated seeding begins at a dominant pinning site and proceeds through sequential residue immobilization (Fig. 1 C), we should observe faster NMR peak volume decay from the early-docking residues.

**Fig. 2.**
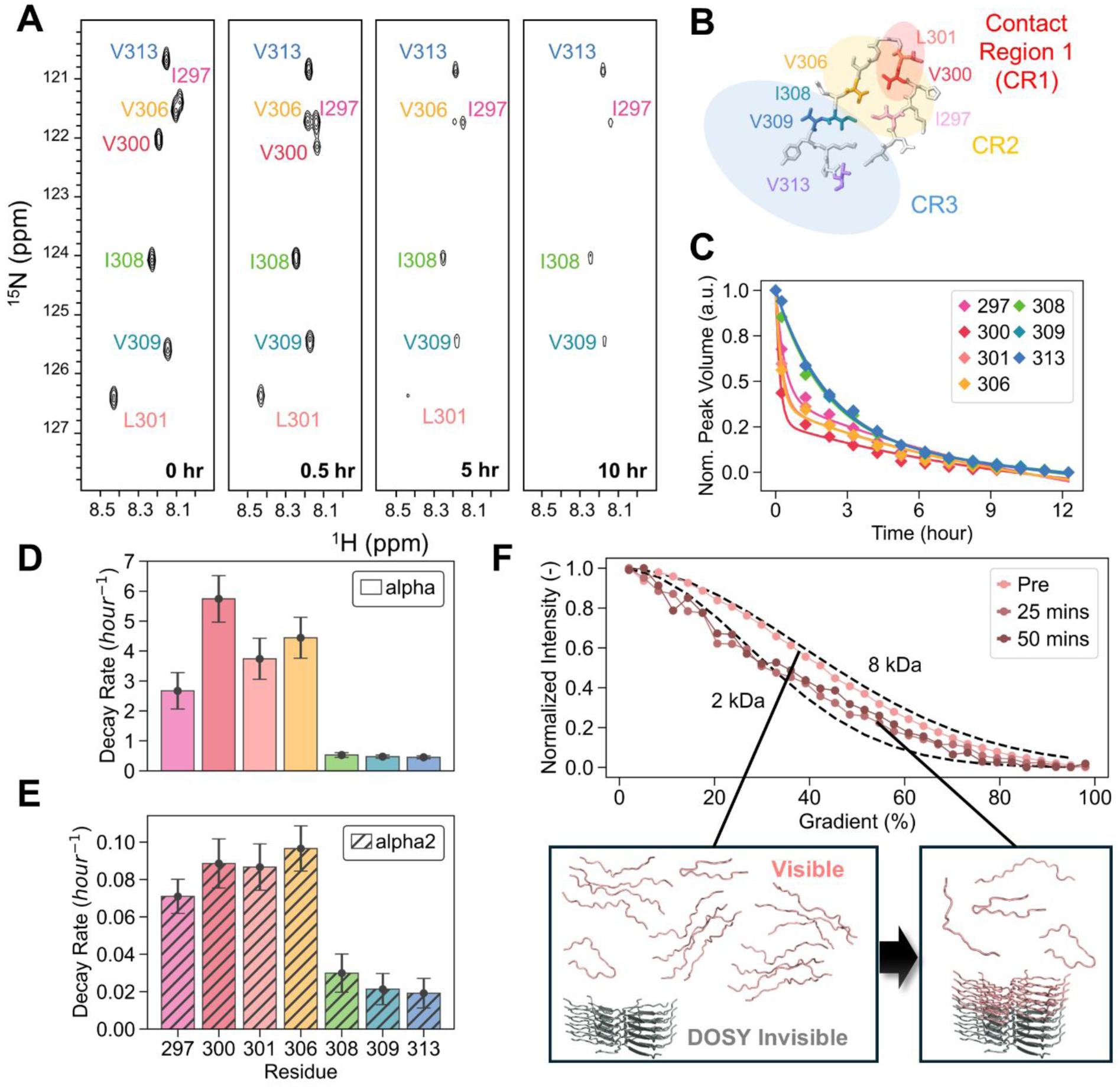
tau jR2R3-P301L peptide templated seeding: (A) Representative ^1^H-^15^N SOFAST-HMQC spectra recorded on jR2R3-P301L templated seeding (298K) induced by a mix of 250 μM ^13^C/^15^N-VLI-jR2R3-P301L with 250 μM jR2R3-P301L fibril seeds over a period of 12 hours. The first spectrum at t = 0 h was recorded before the addition of fibril seeds. (B) The molecular structure of jR2R3-P301L fibril with the sequence of contact regions observed in ^1^H-^15^N SOFAST-HMQC highlighted with different shade colors accordingly (C) Time dependence of the ^1^H-^15^N peak volumes from site I297 (pink), V300 (red), L301 (orange), V306 (yellow), I308 (aqua), V309 (blue), and V313 (purple) on jR2R3-P301L during the templated seeding. The peak volume was obtained from the serially acquired ^1^H-^15^N SOFAST-HMQC spectra and normalized by the maximum changes along the time span of 12 hours. (D) Fast decay rate α and (E) slow decay rate α2 from the biexponential fitting of the peak volume decay in (C). (F) ^1^H DOSY curves before (red) and after the start of templated seeding (light blue, orange), overlapping with the simulated DOSY curves (black dotted) that represent 2 and 8 kDa species. The inlet schematic illustrates the incorporation of oligomeric species as part of the fibril in templated seeding.

A series of ^1^H-^15^N SOFAST-HMQC spectra was recorded after the start of the seeding reaction to track the change of ^1^H and ^15^N spectral properties of seven selected residues I298, V300, L301, V306, I308, V309, and V313 of jR2R3-P301L (Fig. 2 A). The ^1^H-^15^N correlations for these 7 residues have indistinguishable peak intensity and integrated volume before adding the jR2R3-P301L fibril seeds (t = 0 hr, Fig. 2 A). Upon seed addition, the peak volume of these correlations decay at different rates from 0.5 hr to 10 hr upon initiating the templated seeding. 2D contour spectra showed the peak volume of site V300 (red) decayed the fastest (Fig. 2 B, contact region 1, CR1), followed progressively by contact region 2 (CR2) with L301 (light red), V306 (orange), and I297 (pink). The rest of the sites I308 (green), V309 (blue), and V313 (purple) show the slowest decay and were grouped as contact region 3 (CR3). A quantitative comparison of normalized peak volumes from ^1^H-^15^N SOFAST-HMQC spectra confirmed the same trend, with site V300 (red) decaying most rapidly, followed by L301 (light red), V306 (orange), and I297 (pink) in a sequential manner (Fig. 2 C, detailed fitting of normalized peak volumes in Fig. S2).

We next fit the peak volume change using a bi-exponential decay model to compare the peak volume decay rates of selected amino acids (Fig. S2), where α represents the fast decay exponent (Fig. 2 D) and α₂ the slow decay exponent (Fig. 2 E). Within the dock-lock framework of templated amyloid fibril elongation (26), the fast decay rate α reflects the rapid, early docking of residues to the fibril interface, while the slower component α₂ reflects mostly the subsequent, slower disappearance of soluble tau as the fibril grows; we do not assign a unique microscopic mechanism to this slow phase (see Discussion). For both α and α₂, faster rates are found for sites V300, L301, V306, and I297 in CR1 and CR2, suggesting an early engagement of these residues with the fibril surface. Site V300 is clearly a dominant interaction site with the highest decay rates, with α = 5.74 ± 0.78 followed by clear second, third, and fourth values of α = 4.44 ± 0.68 at site 306, α = 3.74 ± 0.68 at site 301 and α = 2.67 ± 0.61 at site 297, while the α values at sites I308, V309 and V313 in CR3 fall below 0.55. In contrast, there is no single dominant site with the highest α₂ values; rather, residues at the bottom of the U-fold (V300, L301, V306, and I297 in CR1/CR2) share similarly elevated α₂ values around 0.086 ± 0.009 relative to the two terminal segments of the U-fold (308, 309, and 313 in CR3, α₂ values around 0.024 ± 0.005). Together, these results demonstrate that templated seeding is initiated through a single dominant interaction hotspot, here centered at V300 in CR1.

Having established that soluble tau species dock onto the fibrillar tau surface, we next decipher whether the soluble tau species engage the fibril surface primarily as a monomer or oligomer. Soluble oligomers may exist in equilibrium with monomers or may form through seed-induced secondary nucleation during templated seeding (26, 36). Although exchange between monomer and oligomer can, in principle, reduce peak intensities through increased transverse relaxation, the fact that normalized peak volumes and intensities decay with the same residue-specific ordering and comparable rates (SI Appendix, Fig. S3 B&C) indicates that no significant exchange-driven line broadening occurs and that the monomer–oligomer equilibrium remains effectively unchanged during seeding. This behavior is consistent with depletion of the soluble population rather than conversion of monomers into new oligomeric species. Still, this observation alone does not distinguish whether monomers are consumed to form fibrils while new soluble oligomers form, or whether pre-existing oligomers are directly depleted. Hence, our next objective is to determine whether oligomer formation precedes, accompanies, or results from the initiation of seed-induced aggregation.

### jR2R3-P301L oligomers are present before fibril seed addition and are rapidly consumed

2D diffusion-ordered spectroscopy (DOSY) measurements were performed to characterize the oligomeric distribution of soluble tau by measuring changes in average diffusivity before and after initiating seed-induced aggregation (37). In DOSY experiments, the diffusivity of soluble tau species is determined from the rate of attenuation of their 1H NMR echo signal with stepwise increasing pulsed field gradient (PFG) amplitudes. Diffusion profiles can be visualized and compared by plotting the normalized integrals of methyl ^1^H resonances from soluble tau species (monomers, and oligomers if present) against a series of PFG amplitudes (38). Greater movement of the ^1^H bearing species across the PFG causes greater dephasing of the ^1^H NMR. Overall, species with a faster diffusivity exhibit greater integral attenuation under identical pulse field gradient strength, whereas more slowly diffusing species lead to moderate attenuation. We refer to the echo amplitude decay with increasing PFG amplitude as “DOSY decay” henceforth. The formation of oligomers in the tau soluble species would result in a more dampened DOSY decay compared to that of monomeric tau.

The DOSY decay of jR2R3-P301L before and 25 min after the start of seed-induced aggregation was compared with two simulated DOSY decays of species corresponding to apparent molecular weights of 2 and 8 kDa, approximating monomeric peptide and small oligomers (e.g. tetramers), respectively (Fig. 2 F, see SI Appendix for details) (39, 40). Before adding fibril seeds, the DOSY decay closely follows the 8 kDa simulated decay, suggesting a heterogeneous mixture between monomers and small oligomers in solution (Fig. 2 F). Within 25 min of seed-induced aggregation, the DOSY decay shifts toward the monomer-like diffusion profile, indicating that the oligomeric population has been selectively depleted while the monomeric species become the dominant visible species. This shift is best explained by rapid incorporation of pre-existing oligomers into larger, NMR-invisible fibrillar assemblies upon binding to the seed surface, leaving monomeric species enriched in solution. It follows that the time-dependent peak decay in the ^1^H-^15^N SOFAST-HMQC series primarily reports on sequential incorporation of soluble tau oligomers onto active fibril ends (schematically shown in Fig 2. F inlet), rather than a monomer-oligomer interconversion driven by secondary nucleation or other mechanisms.

### Enhanced differential hydration around site 300 on jR2R3-P301L fibril surface relative to soluble tau revealed by MD simulation

Next, we examine how the sequential incorporation of soluble tau with active fibril ends relates to its hydration landscape. In solution, the jR2R3-P301L peptide shows increased tetrahedral water structuring and local hydrophobicity around residues 300–301 induced by the P301L mutation (12). We therefore asked whether the fibril surface exhibits a similarly heterogeneous hydration landscape, and whether site 300 in particular harbors hydration water that is susceptible to dewetting and thus capable of driving the initial contact during templated seeding. If hydration water indeed facilitates peptide–fibril association, the NMR-identified interaction hotspot near site 300 should display the most pronounced hydration water change when comparing soluble tau with the fibril surface.

Hydration of the jR2R3-P301L fibril surface was characterized from solvated, equilibrated MD simulations on the top layer in the 3 Å cryo-EM structure of the jR2R3-P301L fibril core (See Materials and Methods for more details), and the resulting per-residue hydration water (black, Fig. 3 A) was subsequently compared with that surrounding the IDP form of jR2R3-P301L (gray, Fig. 3 A). Across the 20 residues from D295 to V314, site 300 showed the largest difference in the number of hydration waters between the disordered and fibrillar states, underscoring the potential role of local hydration changes at this site in facilitating early peptide-fibril contact in templated seeding. The pronounced change suggests that the hydration water remaining at site 300 on the fibril end can be prone to dewetting as an incoming peptide approaches. A comprehensive characterization of the hydration water on the fibril surface is therefore essential for establishing a complete mechanistic picture.

**Fig. 3.**
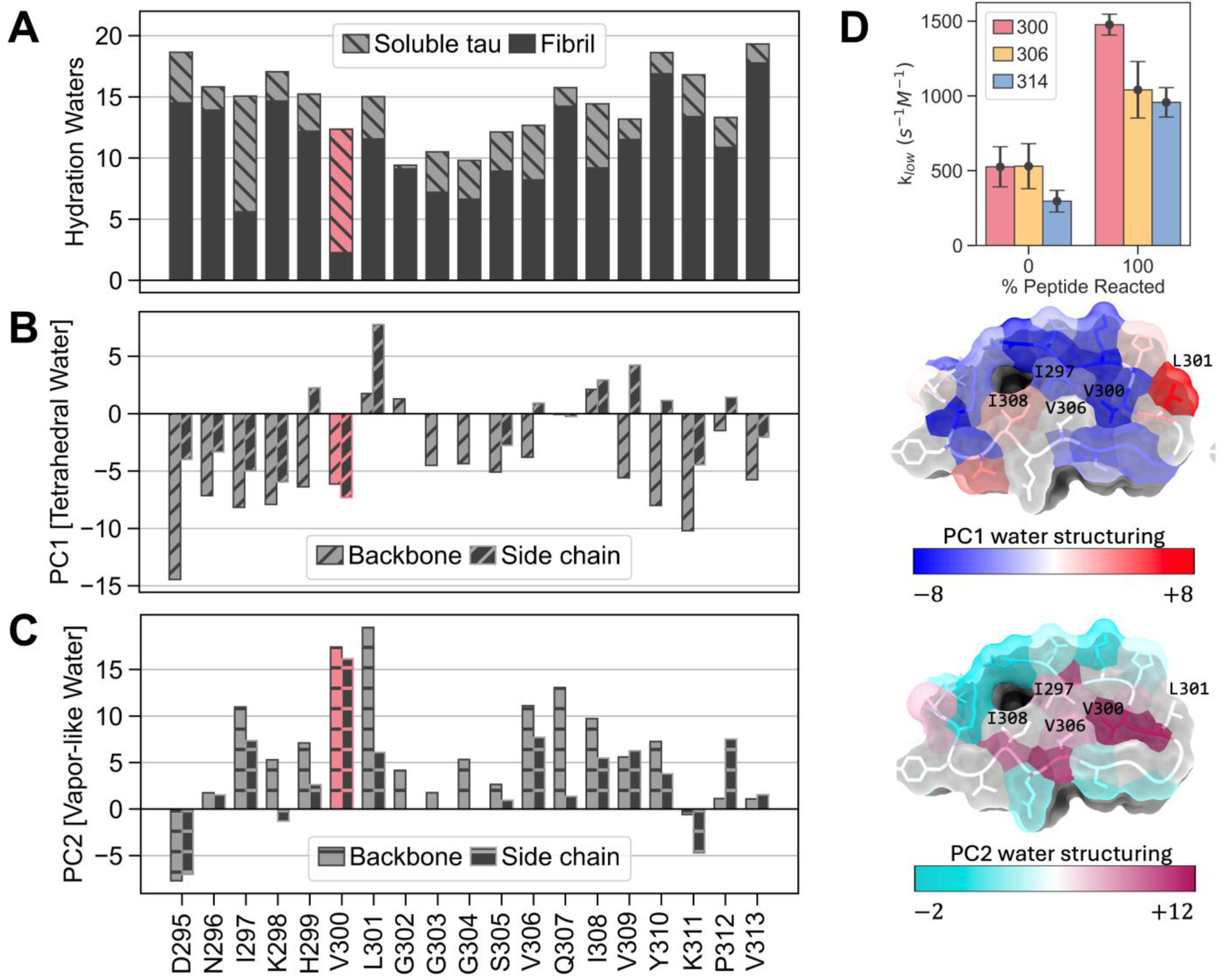
(A) Molecular dynamics simulation of hydration water around soluble tau (gray) and the top layer of the jR2R3-P301L fibril seeds (black). Principal component analysis was performed on the three-body angle distribution of the water hydrogen-bond network at the jR2R3-P301L fibril surface: (B) PC1 reports low-entropy, highly-ordered tetrahedral water population and (C) PC2 reports angularly disordered, ideal-gas-like configuration. For each residue, water structuring around the backbone (grey) and side chain (black) was analyzed separately. The corresponding PC1 and PC2 landscapes are shown next to the bar plots. All the water structuring simulation presented here were carried out using the published jR2R3-P301L fibril structure (PDB: 8V1N) (D) Bound/exchange water dynamics at site 300, 306, and 314 on the surface of jR2R3-P301L fibrils represented by the slow-motion relaxivity k_low_ in ODNP relaxometry measurement.

### Enhanced release-prone hydration water around site 300 on the fibril seed interface

We employed both computational and experimental approaches to characterize the structure and dynamics of water at the templating fibril end. The surface hydration water population can be assessed by its three-body angle distribution, defined by the angle formed between a water molecule and its two nearest neighbors (41). This metric captures the angular organization of the local hydrogen-bond network around the surface. Principal component analysis (PCA) of this angular distribution unveils two dominant modes of water structure (42). The first principal component (PC1) corresponds to a tradeoff between a tetrahedral water population (with ∼109.5° triplet angles) and a simple fluid water population (with ∼60° angles) (43, 44). This tetrahedral population represents a low-entropy state relative to bulk water. Meanwhile, the second principal component (PC2) quantifies a distinct population of water molecules with triplet angles near ∼90°, associated with larger (>1 nm) hydrophobic surfaces and resembling “vapor-like” interfacial water. Surfaces enriched in the tetrahedral population captured by PC1 are expected to be susceptible to dewetting for entropic reasons, as release of these highly ordered waters upon peptide–fibril association increases bulk water’s configurational entropy. In contrast, surfaces enriched in the vapor-like population captured by PC2 are expected to be susceptible to dewetting for enthalpic reasons, since their reduced hydrogen-bond connectivity lowers the energetic cost of water removal. Together, these two metrics provide a quantitative framework for assessing whether the fibril surface presents a hydration environment that can energetically favor tau docking through water release at a specific site (28, 45).

By comparing PC1 and PC2 across all the residues on jR2R3-P301L fibril ends, considering contributions from both backbone and side-chain hydration separately (Fig. 3 B&C), we found that site 301 has the most elevated side-chain PC1 structuring (i.e., more tetrahedral triplet angles) (Fig. 3 B), indicating an enhanced population of low-entropy tetrahedral structural water around this site. Notably, site 300 showed the most elevated PC2 structuring (i.e., more ideal-gas-like structuring) at both its side chain and backbone among all residues, while site 301 was also among the sites with the most elevated backbone PC2 structuring (Fig. 3 C). Together, this combination of elevated PC1 and PC2 structuring suggests that site 300/301 is the region that harbors the most release-prone water, suggesting that it represents a preferred site for initial contact during tau templated seeding via localized hydration-water release at the active fibril surface.

Next, we experimentally probe the local dynamics of hydration water across different regions on the jR2R3-P301L fibril surface using ODNP relaxometry (34). Extensive previous studies have shown a close correlation between surface water dynamics, local water structure, and solvation thermodynamics on accessible surface sites (42, 46, 47). In ODNP, site-specific water dynamics are extracted from ^1^H NMR signal enhancements of water originating from ∼5-8 Å distance of an unpaired electron spin of a nitroxide spin label on the tau surface (34, 48, 49). This spin label can be introduced through site-specific cysteine mutation and labeling, enabling a mapping of hydration along different regions on the fibril surface. The ^1^H NMR signal enhancement profile and *T*_1_ relaxation times are used to derive the value of cross-relaxivity k_σ_ between the electron spin and ^1^H that encodes the translational water diffusivity (on the scale of ∼10 ps) and the value of k_low_ that captures the slow, bound, water dynamics (on the scale of ∼ns) (49, 50). k_low_ reports on the structuring of hydration water on the surface of jR2R3-P301L fibrils and can identify dynamically constrained (bound) water that would entropically stabilize the initial peptide pinning process.

We replaced sites 300, 306, and 314 with a single cysteine, one at a time, for site-specific MTSL labeling. Unlike isotope labeling for NMR, spin labeling is only viable at sites that tolerate a cysteine substitution without compromising the peptide’s ability to dock onto the fibril seed, be templated, and incorporate into fibrils. These three sites lie in the distinct contact regions defined by our time-resolved SOFAST-HMQC data (a rapidly immobilizing region that initiates pinning, CR1; an intermediate-rate association region, CR2; and a slowly associating region, CR3), so this residue-specific strategy probes hydration dynamics at representative sites spanning the hierarchy of sequential association during templated seeding. Each label was positioned on the accessible fibril end by templated seeding of soluble, spin-labeled jR2R3-P301L onto fibril seeds; a seed-to-peptide ratio series then extrapolated the hydration parameters to the condition where unreacted-peptide contributions are minimized and surface-bound labels maximized (fraction of reacted peptide from cw-EPR multicomponent analysis, SI Appendix, Fig. S5; hydration parameters versus seed ratio, Fig. S6; extrapolation to fully reacted peptide, Fig. S7). Among the three sites, klow is highest at site 300 in CR1 (1477 ± 70 s⁻¹M⁻¹), above site 306 in CR2 (1041 ± 189 s⁻¹M⁻¹) and site 314 in CR3 (957 ± 98 s⁻¹M⁻¹), indicating enhanced slow, structured water in CR1 on the fibril surface (Fig. 3 D). The CR1–CR3 difference is statistically significant (p ≈ 2×10⁻⁵), whereas CR2 is intermediate, only marginally distinguishable from CR1 (p ≈ 0.03) and not from CR3 (p ≈ 0.7), so CR1 is the only region with clearly elevated slow structured water. With the water three-body-angle distribution simulations, this identifies a dominant water-pinning site of characteristic structured water around site 300 or 301 in CR1, the same water-pinning site found on the tau monomer.

### Blocking the interaction hotspots on tau fibril seeds reduces their templated seeding efficiency

NMR experiments and MD simulations both identify site300/301 in CR1 as the dominant interaction hotspot for jR2R3-P301L templated seeding. If this hotspot initiates the docking of soluble tau onto the growing fibril end, then blocking it should reduce templated seeding more than blocking other sites. To test this, we spin-labeled soluble jR2R3-P301L with MTSL at sites 300, 306, and 314 – the same sites examined by ODNP. During templated seeding, any spin-labeled peptide that extends the fibril end will position its labels at the fibril surface. Under this condition, the subsequent seeding efficiency provides a direct readout of whether blocking the CR1 hotspot diminishes templated seeding more than blocking other positions.

A ThT fluorescence assay evaluated seeding efficiency when fibril ends were blocked by MTSL labels at different contact regions. Among the three spin-labeled variants, jR2R3-P301L labeled at site 300 in CR1 gave the lowest plateau ThT intensity (Fig. 4 A), indicating the strongest inhibition of templated seeding when CR1 is blocked. Negative-stain TEM collected 12 hr after seeding confirmed this: blocking site 300 greatly suppressed fibril elongation, yielding fibrils (Fig. 4 B) that closely resembled the original seeds (Fig. S1 C) in both length and quantity (fibril-length statistics in Table S2, SI Appendix), whereas spin labels at CR2 or CR3 permitted substantial elongation. As a control that this deficit reflects the fibril hotspot rather than a spin-label-impaired soluble tau, fibril seeds pre-capped with spin-labeled peptide were washed and then reacted with fresh, unlabeled tau peptides; only seeds capped at site 300 showed impeded locking, while all remained active for initial docking (Fig. S8).

**Fig. 4.**
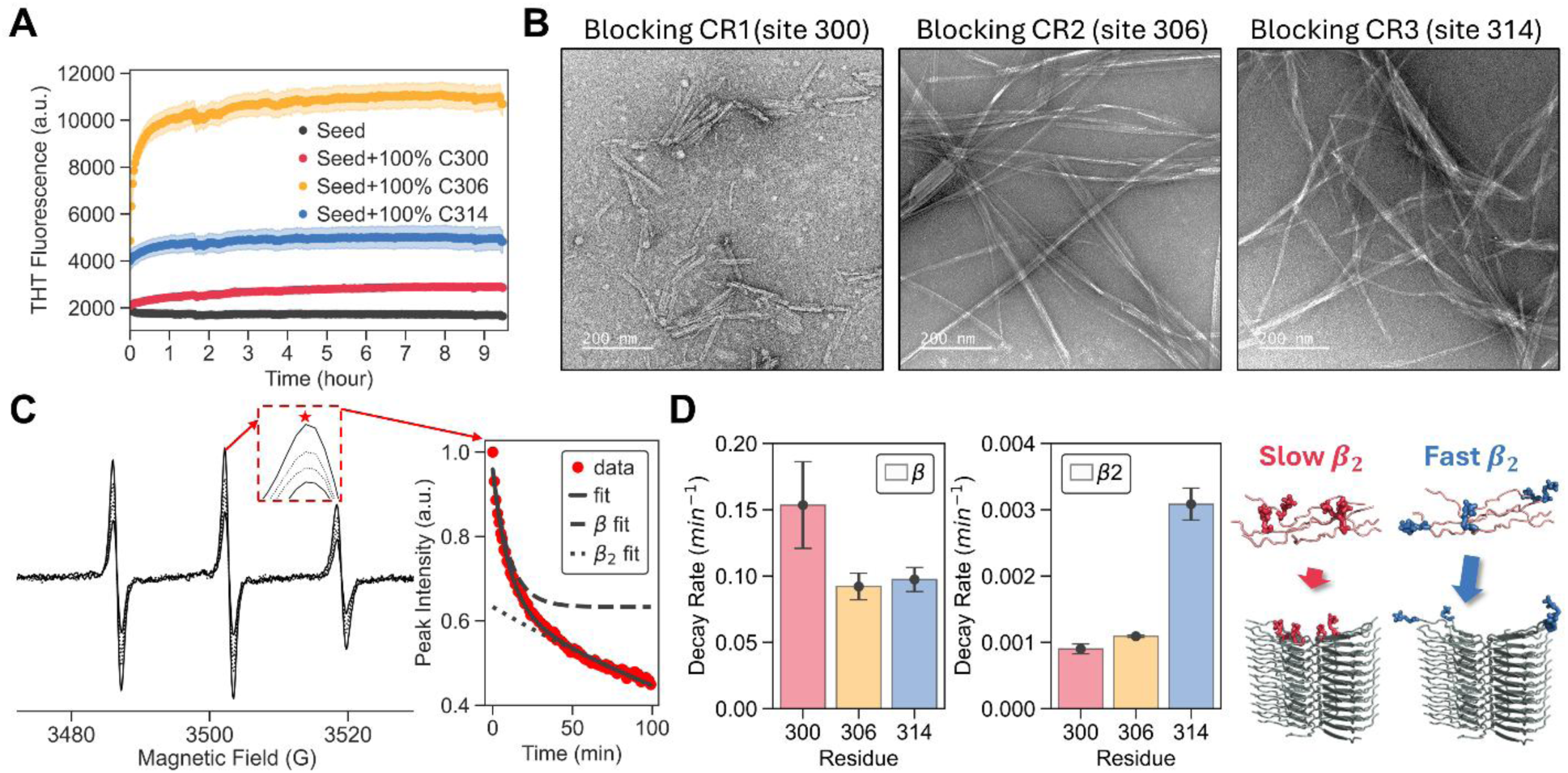
(A) ThT fluorescence of seeding by 100% spin-labeled jR2R3-P301L monomer (100 µM peptide, 100 µM seed) labeled at site 300 in CR1 (red), 306 in CR2 (yellow), and 314 in CR3 (blue), versus seed-only control (black). Labeling site 300 gives the lowest ThT: because 300 is the docking hotspot and the monomer is statistically labeled, a 300-labeled peptide docks but its label caps the fibril end and impedes further locking, whereas labels at 306 and 314 permit continued growth, with 314 locking fastest as the least hindered. (B) nsTEM of the fibrils after the templated seeding of spin-labeled jR2R3-P301L tau. The association rate of spin-labeled jR2R3-P301L to the fibril seeds was tracked by cw EPR spectroscopy: (C) Representative cw EPR spectra collected along the templated seeding are shown, with the decay of central transition intensity was used to quantify the loss of soluble species as they associate with fibril seeds. The decay of the central transition height was fitted with a bi-exponential model, and (D) Fast component β and slow component β₂ from bi-exponential fits of the cw-EPR central-transition decay, for spin-labeled peptide at site 300 in CR1 (red), site 306 in CR2 (yellow), and site 314 in CR3 (blue). The fast component β reports the initial docking before the fibril surface is saturated with spin label and is fastest at site 300; the slow component β₂ reports the subsequent locking after saturation and is slowest at site 300. The inset schematic illustrates the β₂ (locking) step: blocking CR1 (site 300, slow β₂) impedes subsequent association more than blocking CR3 (site 314, fast β₂).

We further examined the docking and locking kinetics of the spin-labeled peptide by cw-EPR. The central-transition height decays as mobile soluble peptide becomes immobilized upon incorporation into the fibril (Fig. 4C; SI Appendix). The decay was fit to a bi-exponential model, yielding a fast component β and a slow component β_2_ (Fig. 4D; triplicate fits shown in Fig. S9). The fast component β reports the initial docking of the spin-labeled peptide at the fibril end and is fastest at site 300 ([1.5±0.1]×10⁻¹ min⁻¹), independently confirming, with a spin probe whose signature is less sensitive to the exchange broadening that can complicate NMR lineshape interpretation, that site 300 docks first. The slow component β_2_ reports the subsequent locking and continued incorporation after the surface is saturated with spin label and is slowest at site 300 ([9.0 ±0.7]×10⁻⁴ min⁻¹) versus site 314 ([3.1±0.2]×10⁻³ min⁻¹). Blocking site 300 thus leaves initial docking intact but kinetically impedes locking and elongation, consistent with the reduced fibril formation seen by ThT and TEM (Fig. 4A, B).

## Discussion

Templated aggregation of jR2R3-P301L on seeding-competent fibrils of the SLS fold characteristic of 4R tauopathies (9, 12) is governed by a single dominant pinning site. We identify it as site 300/301 in CR1 and show that its defining physical signature is locally populated, release-prone hydration water. Soluble tau and the growing fibril end share this site, and localized dehydration there provides the driving force for the initial pinning that enforces in-register stacking into self-amplifying, seed-competent fibrils. Two independent lines of evidence converge on this conclusion: a kinetic one, time-resolved tracking to observe which residue engages the fibril first, and a thermodynamic one, identifying which residue carries the most release-prone water at equilibrium. Both converge on the same site.

Time-resolved ^1^H-^15^N SOFAST-HMQC NMR identified site 300 as the dominant initial pinning contact between soluble tau and the growing fibril end. DOSY showed that jR2R3-P301L exists in solution as small oligomers in equilibrium with monomers before seeding, and that these oligomers dock onto fibril ends and are pulled out of solution, paralleling Aβ₁₋₄₀, where preformed oligomers are consumed early and converted into DOSY-invisible fibrils (38). The SOFAST-HMQC signal decay therefore reports templated association of soluble tau onto fibrils rather than monomer-to-oligomer conversion (36), which is an assignment independently supported by cw-EPR, whose faster initial decay (β) at site 300 marks early association, while cw-EPR lineshape is insensitive to exchange broadening that can complicate NMR lineshape analysis (Fig. 4 D, Fig. S9). Placing these rates in a single dock-lock sequence (Fig. 1D;Fig. 2 D,E), the fast NMR component α (2.7 to 5.7 hr⁻¹, Fig. 2 D) and the fast EPR component β (fastest at site 300, Fig. 4 D, Fig. S9) both report docking on a tens-of-minutes timescale, which is faster than the single-time-constant ThT locking rate (∼0.8 hr⁻¹, Fig. 1 D) and showcases that docking precedes locking. The slow NMR component α₂ (∼0.09 hr⁻¹, Fig. 2 E) is slower than the locking rate, likely not because of a distinct slow chemical step but likely because NMR peak loss is governed by slowed dynamics, through local immobilization and progressively slower tumbling as the assembly grows, rather than β-sheet content; α₂ therefore convolves locking with continued fibril-size growth, while the EPR locking step β₂ (slowest at site 300, Fig. 4 D, Fig. S9) is consistent with this step. Negative-stain TEM confirms fibril elongation within the first ∼10 min (Fig. S1 D), with continued but harder-to-resolve growth thereafter at our high fibril densities, consistent with irreversible, productive locking.

The governing process under our seeded-growth conditions is directed docking-then-locking. Reversible surface exchange, characterized in other amyloid systems by dark-state exchange saturation transfer (DEST) and related methods (Fawzi et al., 2011) describes a steady-state regime (51), including of slow or non-growing assemblies, and likely run in parallel with the productive pathway discussed here; our measurements instead capture net, irreversible growth, underscored by three independent observations. First, negative-stain TEM shows fibril-length extension from seed to fibril within ∼10 min (Fig. S1 D), which surface adsorption alone cannot produce and requires an irreversible locking step. Second, the fast phase α is residue-specific and faster than the ThT rise, identifying an early, site-specific docking step that precedes the cross-β sheet formation measured by ThT fluorescence. Third, the time-dependent ^1^H and ^15^N chemical shifts and linewidths from the same SOFAST-HMQC series (SI Appendix, Fig. S10) show two regimes: nearly all shift and linewidth change occurs in a fast phase (0 to 1 hr), after which these parameters become time-invariant while peak volumes continue to decay, whereas a purely non-specific model in fast exchange would drive ongoing shift and linewidth evolution throughout. Together, these observations are best explained by directed docking at the pinning hotspot followed by locking that extends the fibril, summarized schematically in Fig. S11. Our central conclusion, the residue-resolved identification of a dominant pinning hotspot on the pathway to growth, does not depend on the detailed kinetics of the slow phase α₂. Bulk ThT alone cannot separate one-step from two-step elongation; its initial rate rises monotonically with monomer concentration, as expected when the elongation-competent pool scales with concentration (Fig. S12, detail in SI Appendix) (53). Because ThT reports only locking, it is the residue-resolved NMR and spin-label data that resolve the docking step and localize it to site 300.

These kinetics acquire physical meaning when read against the hydration landscape, because the residue that docks first is the residue that carries the most release-prone water. Extending our previous work showing that the P301L mutation enhances fibril formation by promoting aggregation-prone conformations and enhancing local structured hydration around CR1 in the IDP state (12), we show that the fibril end is pronouncedly dehydrated at site 300 relative to that IDP state. Principal component analysis of local water structuring identifies sites 300 and 301 as carrying the strongest release-prone hydration signature, with the lowest energetic cost for the interfacial water eviction that fibril formation requires, directly supporting water release as the driving force for the pinning event resolved by NMR. Indirect umbrella sampling (INDUS) of surface hydrophobicity, which quantifies how readily hydration water is depleted under increasing bias (SI Appendix, Fig. S13) (52, 53), places CR1 among three hydrophobic patches, but because the metric is highly sensitive to side-chain orientation on the planar fibril surface it cannot by itself resolve a single hotspot; the water-structure analysis does. Hydrophobicity is thus necessary but not sufficient: it is the structure of the interfacial water, not bulk hydrophobicity, that singles out the pinning site. Because pre-existing oligomers are the primary soluble species recruited at the earliest stages of seeding, their hydration properties should also shape which species are most extension-competent; while fibril-end hydration is the dominant determinant of pinning in our system, how hydration varies across soluble oligomeric states is an important direction for future work.

This is the first demonstration that templated tau aggregation is governed by a single dominant pinning site whose defining signature is locally populated, release-prone hydration water that facilitates in-register stacking into seeding-competent fibrils. The critical finding is not the identity of site 300 in jR2R3-P301L but the broader principle that tau templated aggregation requires a dominant pinning site. We expect that site to shift with the tau proteoform under disease-associated mutations and post-translational modifications (54–56) such as phosphorylation (57, 58); our recent study of axial water stabilization in phosphorylated tau suggests surface PTMs could similarly modulate hydration and residue-specific association dynamics, a promising avenue for future investigation (59). That spin-labeling site 300 alone effectively suppressed seeding shows that regions beyond the canonical VQIINK and VQIVYK hexapeptide motifs can govern the initiation of soluble tau-fibril interactions, expanding therapeutic targeting beyond established amyloid cores to the free-energy contribution of hydration-water release from fibril surfaces. Given the heterogeneous hydration landscapes observed across tau fibril polymorphs, including localized water pockets on heparin-induced 0N4R tau fibrils (60), future anti-propagation strategies may exploit dewetting-prone regions as generalizable targets across diverse tauopathies.

## Materials and Methods

### Fibril Seed Preparation

The jR2R3-P301L peptide without isotope enrichment used for fibril seed preparation was purchased from GeneScript. The peptides were dissolved in an ammonium acetate buffer (20mM ammonium acetate, 50 mM NaCl, pH 7.4) supplemented with 5 mM EDTA and 1 mM EGTA to prevent clumping of filaments^1,2^, and then diluted to 50 μM to be mixed with heparin at a 4:1 molar ratio to induce fibril formation. The mixture was shaken continuously at 200 rpm in an incubator at 37 °C for over 24 hours. The formed fibril was concentrated by a 100 kDa cutoff concentrator (Amicon) to less than 100 μL and diluted by MilliQ water to 4 mL at least 4 times to wash away both the excess heparin and Tau peptides that were not incorporated into the fibrils. The fibril was then lyophilized by a FreeZone 2.5L -84 °C Benchtop Freeze Dryer (Labconco) to determine its weight and reconstituted to 500 μM by the desired buffer. The fibril seed was then prepared by sonicating the reconstituted fibril for 1 min using the microtip of a Qsonica sonicator (30% duty cycle, pulse mode, 1 s on-1s off) right before the fibril seeding was monitored by ThT fluorescence assay or spectroscopy tools.

### Thioflavin T (ThT) Fluorescence Assay

All experiments were performed with a Tecan Spark fluorescent plate reader. Each condition was prepared in triplicate with 250 μM ^13^C/^15^N-VLI-jR2R3-P301L and 20 μM ThT in 20 μL ammonium acetate buffer. The seeding fibrillization sample was prepared with 250 μM of seed prepared by the method described above, and the heparin-induced fibrillation was prepared with 62.5 μM heparin. The 100% spin-labeled samples were instead prepared using 100 µM peptide and 100 µM seed. The samples were distributed into a 384-well plate (Corning low-volume non-binding surface black with clear flat bottom) and equilibrated at 25°C. The fluorescence intensity was measured at (excitation=440 nm, emission=485 nm) every 2.5 minutes until a plateau was reached.

### TEM Analysis

All TEM images were acquired by JOEL 1400 TEM in the Northwestern University Atomic and Nanoscale Characterization Experimental Center (NUANCE). 5 µL of fiber sample was applied to a square 200 mesh electron microscopy copper grid with a Formvar-coated carbon film (Electron Microscopy Sciences) that is pre-treated with glow discharge. The grid was blotted dry with filter paper after 60 seconds. Next, 5 µL of 1.5 w/v% uranyl acetate was added to the grid to stain the sample, and the grid was blotted dry after 60 seconds.

### NMR Spectroscopy

All solution NMR experiments were done at 25 ^°^C using a Bruker Neo 600 MHz system with a QCI-F cryoprobe. The peptides with ^13^C/^15^N isotope-enriched on valine, leucine, and isoleucine (^13^C/^15^N-VLI-jR2R3-P301L) were synthesized by the Peptide Synthesis Core at Northwestern University using the solid-phase synthesis method. The peptides were reconstituted to 500 µM in the ammonium acetate buffer with 10 % (v/v) D2O. An equal volume of the reconstituted isotope-enriched peptides and the sonicated fibril seeds was mixed in a 5mm thin-walled precision NMR tube (Wilmad), and the templated seeding was monitored by the time-evolution of two-dimensional (2D) ^1^H-^15^N SOFAST-HMQC spectra up to 15 hours. The total measurement time per 2D spectrum is 16 min with a number of scans (NS) of 16 and a recycle delay of 0.1 seconds.

^1^H-DOSY spectra were acquired with a stimulated-echo pulse sequence using bipolar gradients acquired in 2D mode. The total measurement time is ∼25min per 2D spectrum, with 16 scans in the direct dimension and 32 gradient strengths that were increased linearly from 2 % to 98 % of the maximum value in the z-axis gradient of the probe (48.2 G cm^-1^). To capture the full DOSY curve of jR2R3-P301L monomer and oligomers, the gradient pulse duration was set to 4.0 ms, and the delay between two gradient pulses was optimized to 60 ms. The recycle delay was set to 1s. All spectra were phased and baseline corrected manually by MestReNova (Mestrelab Research). See SI Appendix for further details in data processing.

### Peptide spin labeling

The jR2R3-P301L peptides were ordered from GenScript with site-specific cysteine mutation at sites V300, V306, and an additional site at 314. The peptides were reacted with MTSSL (2,5-dihydro-2,2,5,5-tetramethyl-3-[[(methylsulfonyl)thio]methyl]-1H-pyrrol-1-yloxy, CAS# 81213-52-7) spin labels or dMTSSL (1-Acetyl-2,2,5,5-tetramethyl-Δ3-pyrroline-3-methyl Methanethiosulfonate, CAS# 244641-23-4) diamagnetic spin label analogs at the molar ratio of 5X labels per cysteine. The reactions were incubated at 4°C on a rotisserie shaker and left overnight. The reaction mixtures were then eluted from a PD MIDITRAP G-10 column (Cytiva PN 28918011) using ultrapure water (Invitrogen CAT# 10-977-015). The resulting solution was concentrated to around 0.5-1 mM using nitrogen gas flow and then syringe filtered (Millipore CAT# SLGV004SL) to get rid of any possible aggregates or debris. The resulting spin-labeled peptide concentration was derived using UV-Vis intensity at an absorbance wavelength of 274 nm (Thermo Scientific CAT# ND-ONE-W). The spin label concentration was derived using CW-EPR spin counting, see *CW-EPR* section in SI Appendix for experimental parameters.

### CW-EPR (Continuous Wave Electron Paramagnetic) Spectroscopy

10 µL of samples were loaded into 0.6 mm ID, 0.84 mm OD quartz capillaries (VitroCom 100 mm clear fused quartz, PN: CV6084). CW EPR spectra were acquired using a Bruker EMX X-band spectrometer equipped with a dielectric cavity (Bruker ER 4123D) at around 0.35 T center field, 9.8 GHz microwave frequency, 1.968 mW microwave power, 0.5 G modulation amplitude, 100 kHz modulation frequency, 150 G magnetic field sweep width, 1024 points, and 20.48 ms conversion time and time constant.

### ODNP (Overhauser Dynamic Nuclear Polarization)

The reaction triplicates were prepared at 250 µM spin-labeled peptide with variable seed ratio: 0X, 0.5X (125 µM), 1X (250 µM), and 2X (500 µM). The 0.5X, 1X, and 2X triplicates were incubated at 25°C for 12 hours and then stored at 4°C until measurement. Prior to ODNP measurement, CW-EPR spectrum was taken for each sample, and % unreacted monomer was extracted using the procedures mentioned in SI Appendix. Further details in ODNP data analysis for the hydration variables at the fibril interface were also described in SI Appendix.

ODNP–enhanced ¹H NMR experiments were performed using the X-band EPR hardware setup described in the CW EPR section with additional parts: Bruker cavity (model ER 4119 HS), Bridge12 NMR detection coil (model: B12T ODP 9GHz B12TODP0019 R3) positioned within the cavity, Bruker Avance 300 NMR spectrometer, Bridge12 shim coils (model: E-Shims B12TAES0002) for optimal magnetic field homogeneity, and Bridge12 microwave source (model: MPS 9GHz B12TMPS0006). For the experimental set-up, the NMR coil was tuned to 14.95 ± 0.1 MHz using a vector network analyzer, and the optimal pulse length was determined by a nutation experiment. The automatic data acquisition was implemented using a Python script (package: pyB12MPS^8^), which tells the Bridge12 microwave source to send a trigger pulse to the Bruker NMR console to initiate the Topspin NMR pulse sequence. ¹H signal enhancement curves were acquired by incrementally increasing the microwave power from 0 to 34 dBm over 19 steps and collecting 1D FID. Spin-lattice relaxation times (*T*_1_ and *T*_1,0,0_) were measured using a standard inversion-recovery pulse sequence. The *T*_1_ power curves were collected at 0 dBm, 27dBm, 30dBm, and 34dBm with 15 s of microwave power off before microwave irradiation for each FID to avoid microwave heating. For all the experiments, 15 s microwave irradiation was used to reach saturation of the signal.

### Water Structure Simulation

One strand of the jR2R3 P301L cryo-EM structure (PDB #8V1N) with an internal sidechain zipper was used for the water structure analysis. Residues 312-314 were appended to the structure with Pymol (see jR2R3_P301L_appended.pdb). The structure was solvated, and counterions were added before energy minimization. The structure was simulated for 5 ns in an NPT ensemble with the a99SB-disp force field and with the alpha carbons of residues 295-311 (those resolved with cryo-EM) restrained with a harmonic restraint (1000 kJ/mol spring constant). The 3-body water angle distribution around each sidechain and each backbone was measured, as previously described (12, 41), with a 4.25 Å cutoff used to define hydration waters relative to heavy atoms in the amino acid. Each amino acid’s sidechain or backbone 3-body water angle distribution was subtracted from the bulk water distribution (see bulk_water_triplets.csv), and we took the dot product of the resulting distribution with a principal component distribution (35) (see principalComps.csv) and plotted these values (or colored the structure by these values) in Fig. 3C for PC2.

## Author Contributions

C.H., K.T., S.H. designed research. S.L., S.N., M.S.S., J.E.S. designed simulations. C.H., K.T., S.N., S.L., E.H, K.M. performed research. C.H., K.T, S.L., S.N. analyzed data. C.H. and S.H. wrote the paper.

## Acknowledgments

The study of the seed templated tau aggregation pathway was supported by the National Institutes of Health (NIH) NIA under Grant Number R01AG056058. The study of the minimal prion design and seeding of tau was supported by the Tau Consortium of the Rainwater Charitable Fund. The ODNP study of the role of water in protein interactions was supported by NIH MIRA under Grant Number R35GM136411 and the Deutsche Forschungsgemeinschaft (DFG, German Research Foundation) under Germany’s Excellence Strategy (EXC-2033, project no. 390677874). The W. M. Keck Foundation (www.wmkeck.org) supported the ongoing experimental and computational method developments for the tau shape propagation study. Computational modeling was supported by NSF-ANR MCB/PHY 2423885. Simulations were performed using resources of the Extreme Science and Engineering Discovery Environment, which is supported by the NSF grant ACI-1548562 (project TG MCA05S027 using the Purdue Anvil Cluster and Texas Applied Computing Center Stampede2 Cluster) from the Advanced Cyberinfrastructure Coordination Ecosystem: Services & Support (ACCESS) program, which is supported by U.S. National Science Foundation grants #2138259, #2138286, #2138307, #2137603, and #2138296 and the computational facilities purchased with funds from the National Science Foundation (CNS-1725797) and administered by the Center for Scientific Computing (CSC). The CSC is supported by the California NanoSystems Institute and the Materials Research Science and Engineering Center (NSF DMR 2308708) at UC Santa Barbara. SL and JES acknowledge the generous donation of an anonymous donor, and JES acknowledges support from the NIH MIRA under grant R35GM163771.

^13^C/^15^N isotope-enriched jR2R3-VLI-P301L peptide synthesis was performed at the Peptide Synthesis Core Facility of the Center for Regenerative Nanomedicine at Northwestern University. This facility has current support from the Soft and Hybrid Nanotechnology Experimental (SHyNE) Resource (NSF ECCS-2025633). The Center for Regenerative Nanomedicine, Northwestern University Office for Research, U.S. Army Research Office, and the U.S. Army Medical Research and Materiel Command have also provided funding to develop this facility. NMR, ODNP, and EPR studies were conducted in the Integrated Molecular Structure Education and Research Center (IMSERC) at Northwestern University, which has received support from the Soft and Hybrid Nanotechnology Experimental (SHyNE) Resource (NSF ECCS-2025633), NIH 1S10OD012016-01 / 1S10RR019071-01A1, and Northwestern University. We thank Dr. Yongbo Zhang for his excellent technical assistance. Electron microscopy was conducted at the BioCryo Core within the Northwestern University Atomic and Nanoscale Characterization Experimental (NUANCE) Center, which has received support from the SHyNE Resource (NSF ECCS-2025633), the IIN, and Northwestern’s MRSEC program (NSF DMR-2308691). Last, but not least, we thank Hannah L.H. Weber, Danielle Qin, Victor Zhao, as well as Vishnu Vijayan for their contribution to ThT assay and TEM measurements.

