## Supplemental Information for "A water-pinning hotspot drives templated tau aggregation"

**This PDF file includes:**

Supporting text
Table S1 to S2
Figures S1 to S13

**Other supporting materials for this manuscript include the following:**

### Supporting Information Text

#### $^{1}H$-$^{15}N$ SOFAST-HMQC jR2R3-P301L templated seeding data analysis further details:

The chemical shift assignment of jR2R3-P301L was done by standard triple resonance HNCACB and CBCA(CO)NH experiments on a U-$^{13}C$/$^{15}N$-labeled-jR2R3-P301L peptide. The NMR spectra for assignments were processed using NMRPipe (1) and analyzed with CcpNMR (2) Analysis tools. The assigned $^{1}H$ and $^{15}N$ chemical shifts of the seven selected residues used in the SOFAST-HMQC experiment before and after the seeding are listed in Table S1.

The $^{1}H$-$^{15}N$ SOFAST-HMQC spectra were acquired by using the parameters listed in the Materials and Methods and phased and baseline corrected using the same parameters using NMRPipe. The Fourier-transformed spectra were then fitted by CcpNMR software to get the peak volumes of the seven selected residues before and after the start of the templated-seeding reaction. The peak volumes of different residues were then normalized and fitted by a bi-exponential decay model:

$$I_{normalized}(t) = \frac{I_i(t) - I_i(\infty)}{I_i(0) - I_i(\infty)} = \mu * e^{-\alpha * t} + (1-\mu) * e^{-\alpha 2 * t} + C$$

where $I_i(t)$ is the peak volume at time t in the templated seeding reaction, $I_i(0)$ is the peak volume before the start of the seeding reaction, $I_i(\infty)$ is the peak volume after the reaction reaches a saturation, $\alpha$ is the fast component of the bi-exponential fitting, $\alpha_2$ is the slow component of the fitting, and $\mu$ is the fraction of the fast component. The raw data and the bi-exponential fit of the seven residues are plotted in Fig. S2 A, with the fast decay rate $\alpha$ and the slow decay rate $\alpha_2$ compared in Fig. S2 B & C, respectively.

For each of the seven isotope-enriched residues, the $^{1}H$ and $^{15}N$ amide chemical shifts and linewidths (full width at half maximum, FWHM) were extracted from the time-resolved $^{1}H$-$^{15}N$ SOFAST-HMQC series at t = 0, 1, and 10 hr. Changes were computed separately for each nucleus across two intervals: the Phase 1 change as the 1 hr value minus the 0 hr value, and the Phase 2 change as the 10 hr value minus the 1 hr value. Chemical-shift differences were displayed on a symmetric-logarithmic axis (linear within ±0.005 ppm for $^{1}H$ and ±0.01 ppm for $^{15}N$, logarithmic beyond), and FWHM differences on a linear axis (see Fig. S10).

#### $^{1}H$ DOSY on jR2R3-P301L templated seeding further details:

All $^{1}H$-DOSY spectra were acquired by using the parameters listed in the Materials and Methods and phased and baseline corrected by MestReNova (Mestrelab Research). The methyl peak integrals from jR2R3-P301L were then plotted against different gradient pulse strengths between 2 and 98% of the maximum gradient field strength (48.2 G $cm^{-1}$) to get the DOSY curve. To understand the apparent average MW from the measured DOSY curves, we overlaid the data with two simulated DOSY curves corresponding to jR2R3-P301L monomers (~2kDa) and tetramers (~8kDa). The conversion of apparent MW to diffusivities was done by the empirical relationship between molecular radius of hydration and molecular weight in the literature(3, 4), together with the Stokes-Einstein equation:

$$D = \frac{k_B T}{6\pi\eta R_H}$$

where $k_B$ is the Boltzmann constant, T is temperature (298 K), η is solvent viscosity, $R_H$ is the radius of hydration from the measured molecule, and D is the calculated diffusivity. The estimated diffusities of jR2R3-P301L monomers and tetramers from the corresponding $R_H$ and apparent MW were then applied to the Stejskal-Tanner equation to get the simulated DOSY curve.

$$\frac{I}{I_0} = exp\left[-D \cdot \gamma^2 \cdot g^2 \cdot \delta^2 \cdot \left(\Delta - \frac{\delta}{3}\right)\right]$$

where I is the peak intensity with the pulse field gradient applied, $I_0$ is the peak intensity without the gradient pulses applied, γ is the gyromagnetic ratio of $^1H$, δ is the gradient pulse duration, Δ is the diffusion delay between gradient pulses, and g is the gradient strength as a variable.

**CW-EPR Spin Counting:**
5 scan averages were taken for all samples and 4-OH-TEMPO radicals ((4-hydroxy-2,2,6,6-tetramethylpiperidin-1-yl)oxidanyl, CAS# 2226-96-2). The double integral of the CW-EPR spectra was derived using EasySpin (downloaded from https://easyspin.org) based MATLAB GUI (cwEPR version 3.6.0: downloaded from https://www.mathworks.com/matlabcentral/fileexchange/73292-cwepr). The spin concentrations $C_{SL}$ were calculated from calibration curves derived from double integrals of 4-OH-TEMPO radicals and used in ODNP data processing.

**CW-EPR Spectral Component Analysis:**
CW-EPR spectra were fitted using the MultiComponent software (5) developed by Christian Altenbach (downloaded from https://www.biochemistry.ucla.edu/Faculty/Hubbell/software.html). The CW-EPR fitting was done using previously described protocol (6). For each spin label site, the A, G, and R tenors were first determined by fitting the mobile component, i.e. tau soluble peptide spectrum. Once determined, the A and G tensors were constrained for subsequent fitting and the isotropic R tensor of the 1st (mobile) component was also constrained. The second (immobile) and third (spin-exchanged) components were fitted to spectra of fibrils formed by seeding. The second and third component were fitted with an axially symmetric diffusion tensor (tilt angles: $\alpha_D = 0°, \beta_D = 36°, \gamma_D = 0°$). The third component was also fitted with a Heisenberg spin-exchange frequency ($\omega_{SS} = 140\ MHz$) to account for spin labels stacking. The coefficient $C_{20}$ for orienting potential and axially symmetric R tensors were shared for both the second and third components and allowed to vary. The fitted population ($p_1$) of the mobile component was used to calculate the percentage of unreacted tau peptides and used to normalize the ODNP measurements between seeded peptides of different label sites (c.f. Figure S5).

**ODNP data processing further details:**
All data processing was done using functions from the Python package, DNPLab (downloaded from https://github.com/DNPLab/dnpLab). Each FID was processed in the following sequence: left shift to remove points before the FID, exponential line broadening with a line width of 20 Hz, Fourier transform the data into the frequency dimension, manually phase the NMR spectrum, centering the spectrum to 0 ppm, and integrating the data. For the enhancement curve, the data was normalized by dividing the first point, i.e. the signal intensity at 0 dBm power, and plotted against the microwave power in watts (c.f. Fig. S4 A). The value at 34dBm is taken as the maximum enhancement, $E_{max}$, and normalized between samples based on spin concentration. For the $T_1$ inversion recovery curve, the data were plotted against the delay times and fitted to extract the $T_1$ values (c.f. Fig. S4 B). The ODNP hydration variables were extracted using the "dnp.hydration()" function with the following input variables: enhancement curves, the $T_1$ power curves, $T_{1,0,0}$ value (i.e. the inversion recovery rate of the dMTSSL labeled samples at 0dBm), the spin concentration, magnetic field, "smax_model" as "tethered", and "second_order" interpolation method. The quality of the analysis can be seen by the agreement of the fit of the ODNP saturation curve (c.f. Fig. S4 C).

**ODNP Data Interpretation:**
The ODNP hydration variables are extracted using the three experiments mentioned above and represent key information about hydration dynamics described below (6-11):

1) The [1]H NMR enhancement, $\varepsilon(p)$, depends on the degree of polarization transfer from electron to nuclear spins. $\varepsilon(p)$ can be experimentally derived from the integral of NMR peak of [1]H, $I(p)$, in the presence of microwave power $p$, and the integral of NMR peak of [1]H, $I(0)$, without microwave power using:

$$\varepsilon(p) = \frac{I(p) - I(0)}{I(0)}$$

2) The coupling factor, $\xi$, is a unitless variable, which denotes the local efficiency of the ODNP polarization transfer between the free electron and [1]H and only depends on the motion of water within 5-10 Å of the spin label. The coupling factor is a ratio of two fundamental proton relaxivity parameters: $\xi = k_\sigma / k_\rho$.

3) The local cross-relaxivity, $k_\sigma$ $(s^{-1}M^{-1})$, is driven by the electron spin flip excitation close to or faster than the time scale of the electron spin frequency of ~9.8 GHz; therefore, it is selectively sensitive to fast-moving ($ps$ scale), loosely bound water. $k_\sigma$ is experimentally derived from enhancement, $\varepsilon(p)$, and spin-lattice relaxation time, $T_1(p)$, at varying microwave powers using:

$$k_\sigma s(p) = \left(\frac{|\omega_H/\omega_e|}{C_{SL}}\right)\varepsilon(p)T_1^{-1}(p)$$

where, $C_{SL}$ is the spin concentration, $\omega_H$ and $\omega_e$ are the Lamour frequencies of proton and electrons, $s(p)$ is the saturation factor, which quantifies the saturation of the electron spin transition across the three hyperfine components of the nitroxide radical at microwave power, $p$. Without microwave power, $s(p = 0) = 0$, and with increasing microwave power, $s(p)$ asymptotically approaches $s_{max}$, which can range from $\frac{1}{3}$ to 1 for nitroxides. To extract $k_\sigma$, the $k_\sigma s(p)$ are fitted to the infinite microwave power using the following equation:

$$k_\sigma s(p) = \frac{k_\sigma s_{max} p}{p_{1/2} + p}$$

Where $p_{1/2}$ is the microwave power at which half of the electron spin transition is saturated and the $k_\sigma s(p) \to k_\sigma s_{max}$ at infinite microwave power. Since $s_{max} = 1$ is a good approximation for biological samples with tethered spin labels, $k_\sigma = k_\sigma s_{max}$. See Fig. S4 C for ODNP saturation curve and the asymptotic fit to extract $k_\sigma$.

4) The local self-relaxivity, $k_\rho$ $(s^{-1}M^{-1})$, includes also the contribution from the fluctuations at the time scale of [1]H Lamour frequency of ~14.8 MHz. $k_\rho$ is experimentally derived by saturation recovery experiment at 0 dBm power to extract spin lattice relaxation time of proton with spin labels ($T_{1,0}$) and without paramagnetic spin labels ($T_{1,0,0}$). With known spin label concentration, $C_{SL}$, $k_\rho$ is calculated using:

$$k_\rho = \frac{1}{C_{SL}}\left(\frac{1}{T_{1,0}} - \frac{1}{T_{1,0,0}}\right)$$

5) The relaxivity, $k_{low}$ $(s^{-1}M^{-1})$, is driven by the nuclear spin single-flip excitation close to or faster than the time scale of [1]H Lamour frequency of ~14.8 MHz; therefore, it is selectively sensitive to the slow-moving ($ns$ scale), tightly bound water. $k_{low}$ can be determined from $k_\rho$ and $k_\sigma$ using:

$$k_{low} = \frac{5}{3}k_{\rho} - \frac{7}{3}k_{\sigma}$$

6) The translational correlation time, $\tau_c\ (s)$, denotes the lifetime of the dipolar interaction between the electron and proton, i.e how long a proton molecule stays within the interaction distance of the spin label. $\tau_c$ can be extracted from the coupling factor, $\xi$, by analytically translating $\xi$ using the force-free hard-sphere model, FFHS. The FFHS method assumes the translational diffusion dominates the dipolar coupling between electron and nuclear spins, with no rotational diffusion. The FFHS spectral density function is the following:

$$\xi(B_0; \tau_c) = \frac{6J(\omega_e + \omega_H; \tau_c) - J(\omega_e - \omega_H; \tau_c)}{6J(\omega_e + \omega_H; \tau_c) + 3J(\omega_H; \tau_c) + J(\omega_e - \omega_H; \tau_c)}$$

7) The local diffusivity, $D_{local}\ (m^2s^{-1})$, denotes the diffusivity of hydration water within 5-10 Å of the spin label. $D_{local}$ can be calculated from $\tau_c$ using the following:

$$\tau_c = \frac{d^2}{D_{local} + D_{SL}}$$

where $D_{SL} = 4.1 \times 10^{-9}\ m^2s^{-1}$ is the diffusivity of TEMPO spin probe and $d$ is the distance of approach between the water and the spin probe.

The above ODNP hydration variables from jR2R3-P301L with spin labels on site 300 (red), 306 (orange), 314 (blue) under various seed-to-peptide ratios (0, 0.5X, 1X, and 2X) are compared in Fig. S6, where the 0X condition represents the ODNP hydration variables measured from jR2R3-P301L peptides in the solution and the 2X represents more of the ODNP hydration variables To limit the contribution of unreacted peptides to the hydration dynamics and to maximize the contributions from the spin labels on the fibril surface, the hydration variables ($k_{\sigma}, k_{low}, and\ D_{local}$) derived from the reaction triplicates at varying seed concentration (0, 0.5X, 1X, and 2X) was plotted against the % peptide reacted extracted from the cw-EPR lineshape (Fig. S7 A-C), and a linear fit was applied to extrapolate the hydration variables ($k_{\sigma}, k_{low}, and\ D_{local}$) under the condition of 100% reaction yield (i.e all spin-labeled peptides incorporated to the surface of the fibril seed). The extrapolated hydration variables that represents the hydration on the surface of fibrils were compared to the values from soluble peptides (Fig. S7. D-F), with the error being propagated as:

$$\sigma_{100} = \sqrt{(100 \times \sigma_m)^2 + \sigma_c^2 + 2 \times 100 \times \rho \times \sigma_m \times \sigma_c}$$

Where $\sigma_m$ and $\sigma_c$ are fitting errors of the variables $m$ and $c$, and $\rho$ is the correlation coefficient between $m$ and $c$. (12)

Pairwise differences in extrapolated $k_{low}$ were evaluated by two-tailed *z*-tests ($z = \Delta/\sqrt{(\sigma_A{}^2 + \sigma_B{}^2)};\ p = 2(1 - \Phi(|z|))$ ), using the $1\sigma$ standard errors from least-squares fitting and extrapolation as the parameter uncertainties.

We aim to compare the differences in structured water population across fibril surface sites by examining the extrapolated values of the hydration variables at 0% free peptide. Among the three main hydration variables, we predominantly consider $k_{low}$, which is derived from the relationship of $\frac{5}{3}k_{\rho} - \frac{7}{3}k_{\sigma}$. This choice is justified due to the following:

1) From the conceptual perspective, our objective is to probe the population of low-entropy tetrahedral structural water at each contact regions on the fibril surfaces (site 300 for CR1, site 306 for CR2, and site 314 for CR3) as represented by PC1 in MD simulation (c.f Figure

3B). This energetically favorable water corresponds to the slow-moving water that is susceptible to dewetting and subsequently initiate tau peptide docking onto certain sites. Compared to $k_\sigma$ ($ps$ scale), $k_{low}$ better represents slow-moving ($ns$ scale) water on the fibril surface that is most relevant to the proposed docking mechanism.

2) From the technical perspective, $k_{low}$ provides more robust comparison of water dynamics in the fibril system than $k_\sigma$ or $D_{local}$. $k_\sigma$ depends on fitting the ODNP saturation curve which becomes increasingly unreliable as the system slows or as a large fraction of spin labels becomes buried (conditions that occur at 100% reacted peptide, where most spin-labeled peptides are on the fibril surfaces or within the fibril layers). Similarly, $D_{local}$ depends on the coupling factor, $\xi$, defined as the ratio of $k_\sigma/k_\rho$, and is thus strongly influded by uncertainties in $k_\sigma$. In contrast, across residues 300, 306 and 314, $k_\sigma$ is small in magnitude such that their $k_{low}$ trends with $k_\rho$ (c.f. Figure S6C vs S6D). Experimentally, $k_\rho$ depends primarily on the proton spin lattice relaxation time measured with and without paramagnetic spin labels and is therefore less sensitive to slowed molecular dynamics or buried spin labels. Consequently, $k_{low}$ more reliably reflects the population of slow-moving, structured surface water population.

**CW-EPR Monitored Seeding Reaction:**
250 µM of peptide and 250 µM of sonicated seed were quickly mixed by aspirating and loaded into EPR capillaries. A pseudo 2D EPR spectrum was collected, where each slice is a set of scans. For a 100% spin-labeled peptide reaction, each slice contains 2 scans, and each slice represents 1-minute reaction interval. The maximum of the central nitroxide peak was used to track the amount of spin-labeled jR2R3-P301L soluble species (c.f. Fig. 4 C). The resulting reaction curve was fitted with bi-exponential decay: $\mu * e^{-\beta * x} + (1-\mu) * e^{-\beta 2 * x} + C$ to extract reaction rate (c.f. Figure S9). The intensity of the last time point was divided by that of the first time point to extract the relative reaction efficiency.

**Preparation of pre-capped fibril seed with spin-labeled peptides:**
Pre-capped fibril seeds were prepared by incubating 100 µM seeds with 100 µM MTSL-labeled jR2R3-P301L (site 300, 306, or 314) for 60 min, followed by two rounds of 10 kDa spin-column filtration (9000 g, 7 min) to remove unbound labeled peptide and retain only firmly bound end-capping fibril seeds. The resulting pre-treated seeds were then mixed with 100 µM fresh, unlabeled jR2R3-P301L peptide, and elongation was monitored by ThT fluorescence.

**INDUS**

The equilibrium and Indirect Umbrella Sampling (INDUS) simulations (13, 14) were performed using GROMACS version 2016.3 (15, 16), which was modified to incorporate biasing potentials on the coarse-grained number of water molecules, $N_v$, within a defined hydration volume. This customized version of GROMACS was obtained from the Patel group at the University of Pennsylvania.

All simulations employed the adisp99sb force field (17). Lennard-Jones and short-range electrostatic interactions were truncated at 1.0 nm, while long-range electrostatics were calculated using the particle mesh Ewald (PME) method (18). Protein bonds involving hydrogen atoms were constrained with the LINCS algorithm, and water geometries were maintained using SETTLE (19). The equations of motion were integrated using the Leapfrog integrator with a 2 fs time step.
Protein systems were solvated in a cubic box with at least 1.0 nm between any protein atom and the box edge. Solvation was performed using genbox, and sodium or chloride counterions were added to neutralize the system. We impose position restraints (with strength 1000 $kJ \cdot mol^{-1} \cdot nm^{-2}$) on heavy atoms of the fibers. Energy minimization was carried out using the steepest descent algorithm, followed by equilibration at 300 K and 1 bar. Equilibration consisted of a 100 ps NVT simulation using the Berendsen thermostat, followed by a 100 ps NPT simulation with the Berendsen barostat (20) and the stochastic velocity-rescale thermostat (21). Production simulations were then run for 3 ns for both bound and unbound systems. During production,

pressure was maintained at 1 bar using the Parrinello-Rahman barostat (τ = 1 ps), and temperature was controlled at 300 K using the stochastic velocity-rescale thermostat (τ = 0.5 ps).

To evaluate fiber surface hydrophobicity and construct a hydrophobicity scale for fiber surface residues, the interactions between the top peptide of the fiber and its surrounding hydration water were perturbed using an external linear biasing potential. The peptide hydration volume (probe volume) was defined as the union of spherical sub-volumes with radius $R_v$, centered on each heavy atom—nitrogen (N), oxygen (O), carbon (C), and sulfur (S)—of the peptide's amino acids. Here, $R_v$ = 0.55 nm was chosen to capture the first two layers of hydration waters. This bias imposed an energetic penalty for each water molecule residing within the probe volume (the hydration shell). The INDUS method was employed to modify the Hamiltonian with a bias linear in the number of coarse-grained waters within the probe volume:

$$H_\varphi = H_0 + \varphi N_v$$

where $H_0$ is the unbiased Hamiltonian, φ is a tunable parameter controlling the strength of the bias, and $N_v$ is the coarse-grained number of water oxygens in the probe.

Hydrophobicity at the atomic level was quantified by calculating the normalized hydration level of each protein surface heavy atom, defined as:

$$\langle \rho_i \rangle_\varphi = \langle n_i \rangle_\varphi \, / \, \langle n_i \rangle_0, \quad \text{at } \varphi = 7.0$$

Roughly around this bias strength ($6 < \varphi < 8$) for most proteins, water susceptibility reaches a maximum, resulting in the depletion of hydration water from hydrophobic domains and thus identifying hydrophobic "hotspots" (22). Here, $\langle n_i \rangle_\varphi$ represents the average number of water molecules in a spherical sub-volume centered on atom i in the biased ensemble, normalized by the corresponding hydration in the unbiased ensemble, $\langle n_i \rangle_0$, to account for differences in solvent accessibility.

Fibril surface hydrophobicity was quantified by the ratio of the number of hydration water molecules at maximum susceptibility to the number of hydration water molecules at equilibrium for each residue. The resulting hydrophobicity map revealed a heterogeneous hydration affinity landscape across the cryo-EM-derived fibril surface (23), indicating that different surface regions exhibit distinct propensities for water association and exclusion (Fig. S13). Three distinct hydrophobic patches emerged: a localized region around 301L, a broader patch spanning I297, V300, V306, and I308, and a more pronounced hydrophobic region centered around V309, Y310, and P312. Yet, upon screening alternative side-chain conformations by restraining the backbone heavy atoms while allowing side-chain reorientation, we noticed that the relative hydrophobicity scale in INDUS showed substantial dependence on side-chain orientation on the planar fibril surface in MD simulation, making it difficult to pinpoint a dominant hotspot on the fibril surface using this approach. Overall, the water structure analysis captures angular ordering within the hydrogen-bond network formed by surface water at the interface, and ODNP relaxometry reports hydration dynamics highly correlated with water structure, both of which are more directly linked to the kinetic susceptibility of water release, while the INDUS hydrophobicity map may still provide insights to secondary hydrophobic domains that are thermodynamically preferred for β-sheet stabilization.

**ThT Concentration Series:**

To complement the high-seed conditions (1 to 1 seed to peptide ratio) used throughout this work, we performed a ThT seeded-aggregation series in which the jR2R3-P301L peptide concentration was varied from 50 to 250 μM at a fixed seed concentration of 100 μM (Fig. S12). These conditions were initially omitted because our assays were designed with seeds in large excess over peptide, isolating the docking step under conditions where peptide supply is not the principal limiting factor; the concentration series instead probes how the bulk kinetics respond as peptide availability is varied. Each trace was fit to the lock-limited single-exponential form of the consecutive dock-and-lock scheme (Esler et al., 2000) (24), valid in the limit where surface association is fast relative to conformational locking. The apparent single-exponential ThT rate increased with peptide concentration; because a unimolecular locking step is concentration-independent, this peptide dependence indicates that the observed ThT rate reflects the population of the docked complex that feeds the locking step rather than any change in the intrinsic locking rate.

To characterize this dependence with a model-independent observable, we additionally applied the initial-rate elongation analysis using protocol Step 37 of Meisl et al. (2016) (25). The initial elongation rate increased linearly with peptide concentration over 100–250 μM (through-origin fit, $R^2 = 0.998$), yielding an apparent elongation rate constant $k\_app = 2k_+P_0 = 9.30$ a.u.·μM$^{-1}$·hr$^{-1}$ (Fig. S12); the 50 μM condition was excluded because its initial-rate signal fell within instrumental baseline drift over the chosen window. This linear, non-saturating dependence, together with the absence of a lag phase (Fig. 1D), is consistent with peptide-limited elongation at the seed ends. We did not use the plateau ThT amplitude as a quantitative readout, since at high signal it can become limited by dye availability rather than fibril mass; the initial-rate analysis avoids this by sampling the early regime where ThT tracks β-sheet mass accumulation. While these bulk kinetics are fully compatible with a two-step mechanism, they do not by themselves distinguish dock–lock from single-step templated elongation; the decisive evidence comes from the residue-resolved NMR and spin-label data. The absolute elongation rate constant $k_+$ was not extracted, as neither $P_0$, the number concentration of growth-competent fibril ends in the sonicated seed stock, nor the ThT-to-mass conversion was independently determined.

**TEM Fibril Length Analysis:**

Fibril length distributions for sonicated seed and the C300, C306, and C314 elongation reactions, measured from negative-stain TEM micrographs (50,000×). Fibril contour lengths were traced manually in Fiji/ImageJ (Schindelin et al., 2012) using the segmented-line tool on a single representative micrograph per condition (26), with the spatial scale calibrated to the burned-in scale bar (915 px = 200 nm; 0.2186 nm/px). Table S2 shows median, mean and ± SD of the fibril lengths (n = 34, 33, 21, and 22 fibrils for seed, C300, C306, and C314, respectively).

**Table S1.** 1H/15N backbone resonance assignments for 13C/15N-labeled jR2R3-P301L

| Residue | $^{1}H$ (ppm) | $^{15}N$ (ppm) |
|---|---|---|
| **Before Seeding** | | |
| 297 | 8.079 | 121.40 |
| 300 | 8.191 | 122.03 |
| 301 | 8.428 | 126.42 |
| 306 | 8.101 | 121.54 |
| 308 | 8.227 | 124.08 |
| 309 | 8.143 | 125.60 |
| 313 | 8.150 | 120.69 |
| **After Seeding** | | |
| 297 | 8.140 | 121.74 |
| 300 | 8.135 | 122.16 |
| 301 | 8.431 | 126.38 |
| 306 | 8.183 | 121.73 |
| 308 | 8.245 | 124.03 |
| 309 | 8.173 | 125.44 |
| 313 | 8.178 | 120.87 |

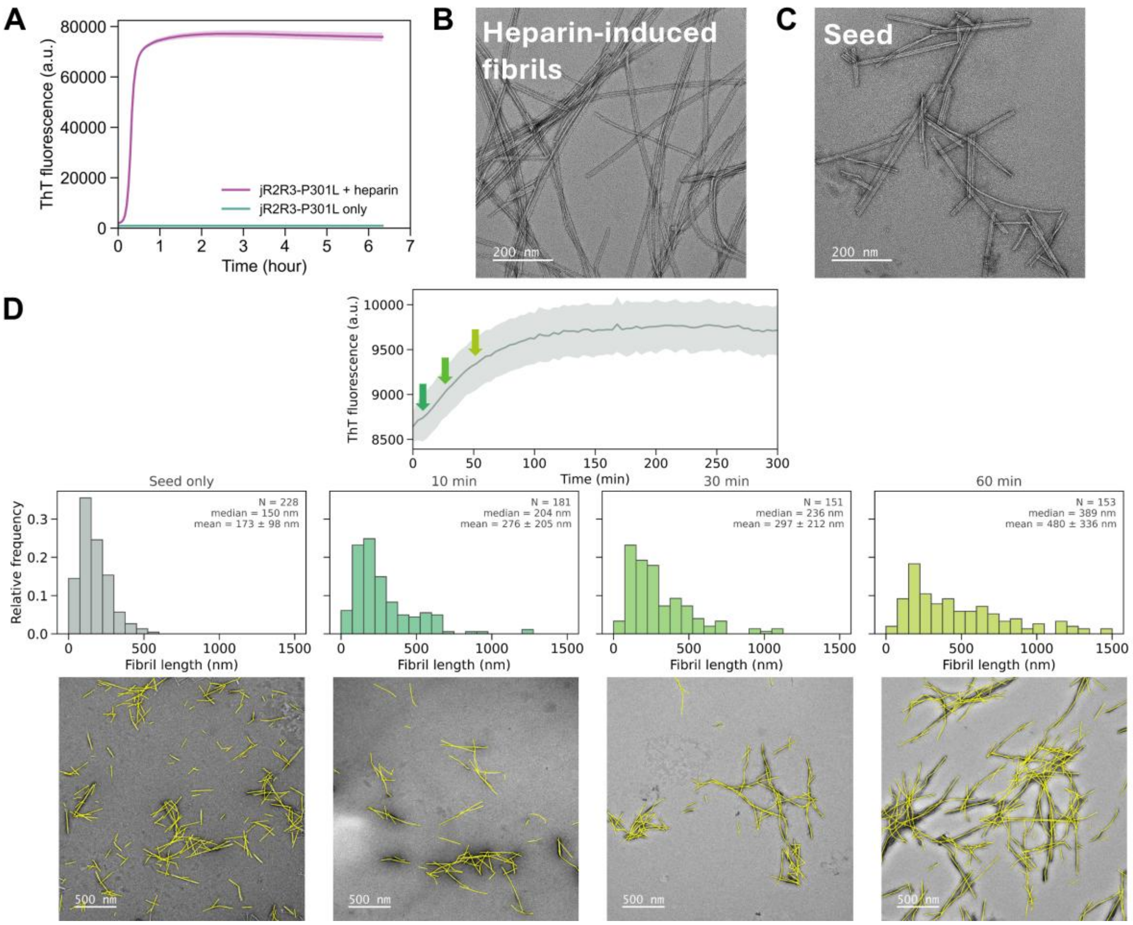


**Fig. S1. Seeded aggregation.** jR2R3-P301L fibril seed preparation. (A) ThT fluorescence of jR2R3-P301L aggregation (blue) induced by mixing the peptide with heparin at a 4:1 molar ratio. The control sample with peptide only (red) shows no self-aggregation in the detected time frame. Representative nsTEM images of (B) the heparin-induced jR2R3-P301L fibrils and (C) the fibril seed prepared by sonicating the heparin-induced fibrils to increase active fibril ends for better templated seeding activities. The fibril end concentration $P_0$ in the sonicated seed stock was not independently determined. (D) Negative-stain TEM of the seeded reaction (250 μM tau peptide, 250 μM seed) at t = 0, 10  min, 30 min and 60 min, showing fibril elongation already evident within the first ~10 min, consistent with the ThT locking rate in Fig. 1 D.

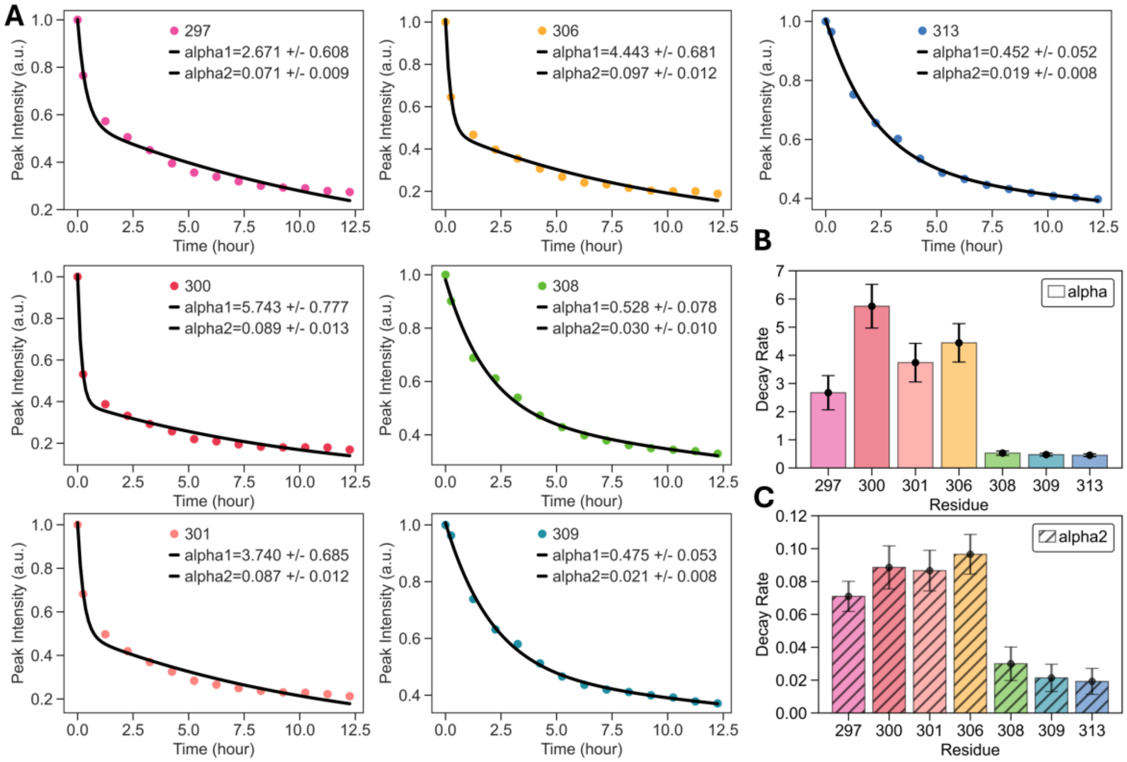


**Fig. S2.** (A) $^{1}H$-$^{15}N$ SOFAST-HMQC time-dependent peak-volume of site I297 (pink), V300 (red), L301 (light red), V306 (orange), I308 (green), V309 (blue), and V313 (purple) fitted with bi-exponential decay: $\mu * e^{-\alpha * t} + (1-\mu) * e^{-\alpha 2 * t} + C$ (B) $\alpha$ comparison between residues. (C) $\alpha 2$ comparison between residues. The error bars in (B) and (C) are reported from the standard error of the curve.

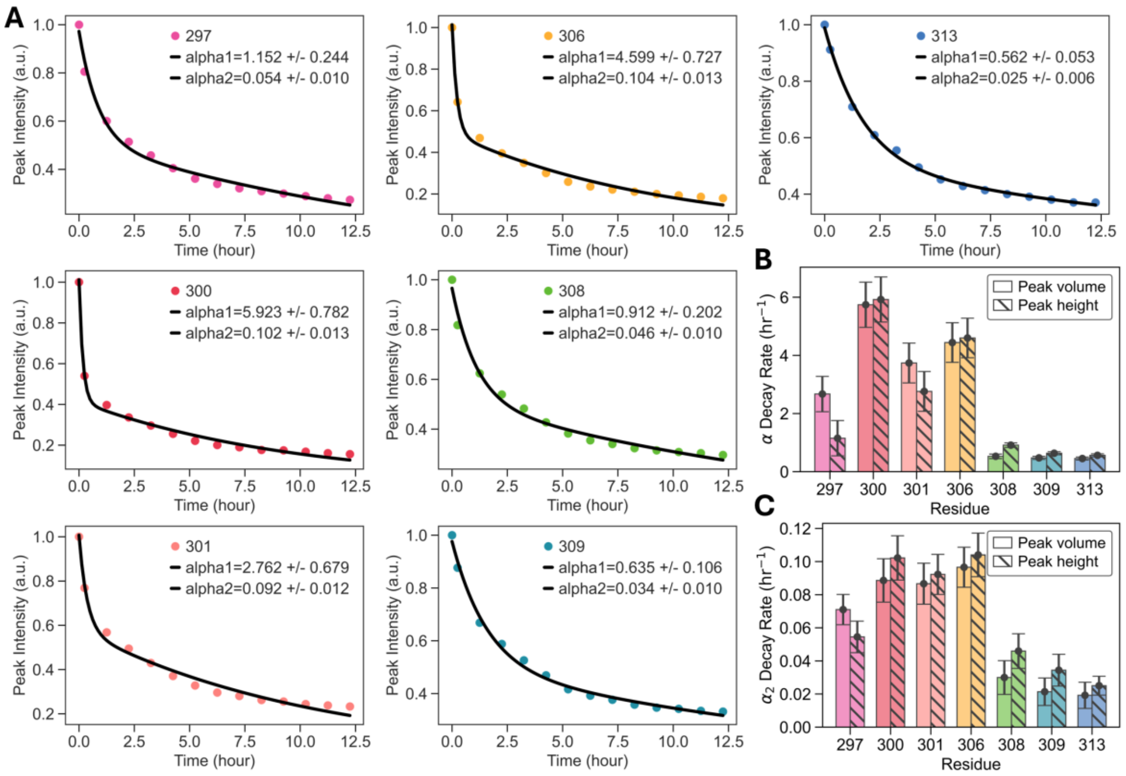


**Fig. S3. Comparison of decay rate constant extracted from peak-volume vs peak-height.** (A) $^{1}H$-$^{15}N$ SOFAST-HMQC time-dependent peak-height of site I297 (pink), V300 (red), L301 (light red), V306 (orange), I308 (green), V309 (blue), and V313 (purple) fitted with bi-exponential decay: $\mu * e^{-\alpha * t} + (1-\mu) * e^{-\alpha 2 * t} + C$ (B) $\alpha$ comparison across residues between peak-volume (solid color) and peak-height (diagonal hatch). (C) $\alpha 2$ comparison across residues between peak-volume (solid color) and peak-height (diagonal hatch). The error bars in (B) and (C) are reported from the standard error of the curve.

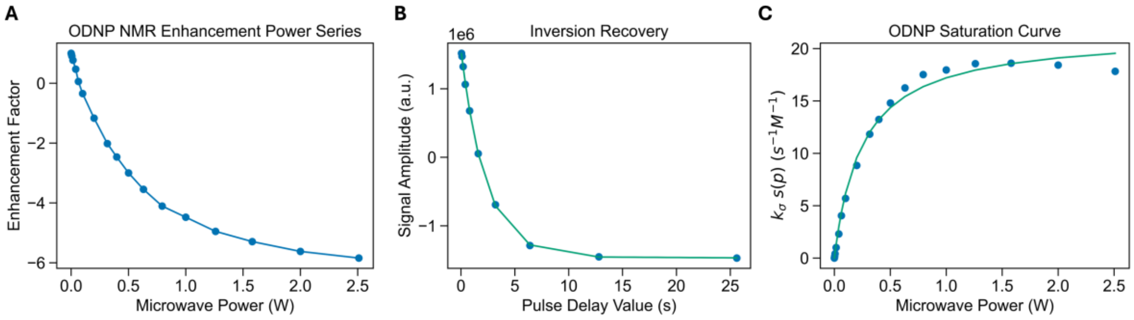


**Fig. S4.** ODNP data processing workflow example. (A) Enhancement vs microwave power. (B) $^{1}H$ NMR peak volume vs pulse delay time. (C) ODNP Saturation Curve.

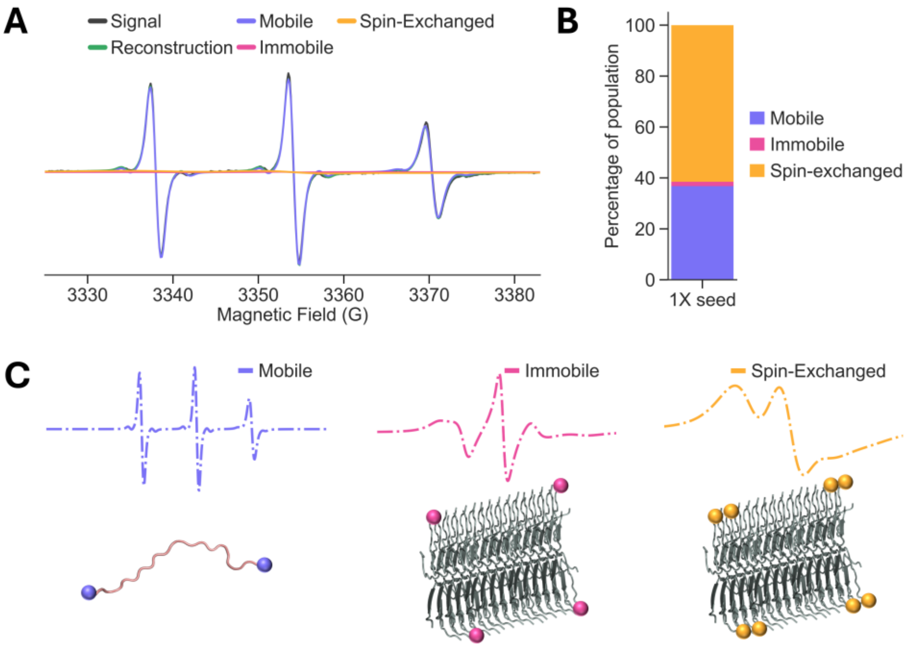


**Fig. S5.** cw-EPR multicomponent fitting to obtain the ratio between the mobile component from soluble species and the immobile component from the spin-labeled jR2R3-P301L peptides incorporated into fibrils in templated seeding. The percentage of the immobile component is interpreted as the extent of reaction and used in the subsequent ODNP data interpretation to get hydration variables on the surface of fibrils. (A) The raw data overlaid with fitting components using experimental data of C300 1X seed reaction as example. (B) The percentage of each population. (C) Zoomed out from (A) of mobile, immobile, and spin-exchanged species with schematic demonstrating each component.

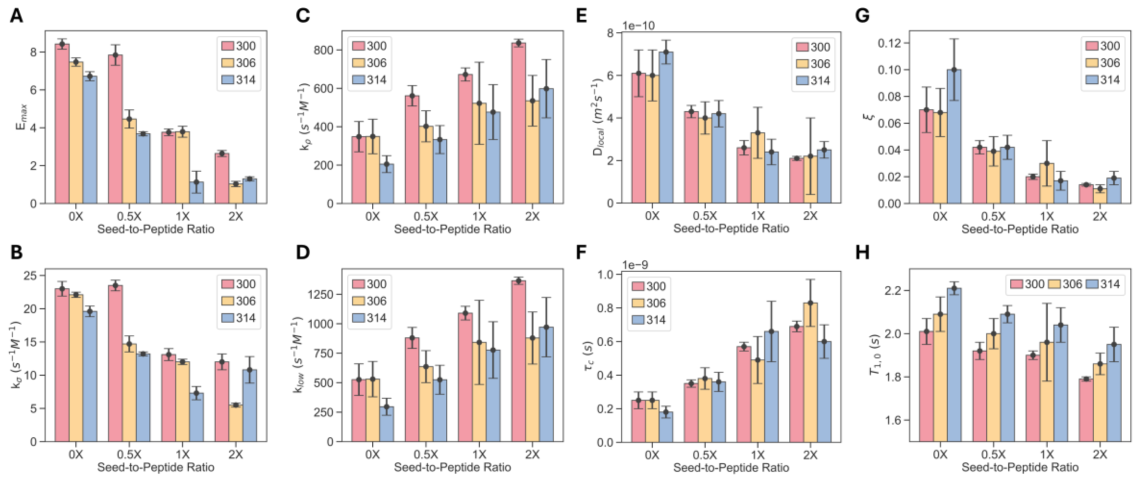


**Fig. S6.** Hydration variables by seed-to-peptide ratio for spin labels on residues 300, 306, and 314. (A) Ehancement, $E_{max}$. (B) Cross relaxivity, $k_{\sigma}$. (C) Self relaxivity, $k_{\rho}$. (D) $k_{low}$. (E) Local diffusivity, $D_{local}$. (F) Correlation time. $\tau_{corr}$. (G) Coupling factor, $\xi$. (H) Relaxation time, $T_{1,0}$. The error bars are reported as 1 standard deviation of the triplicates.

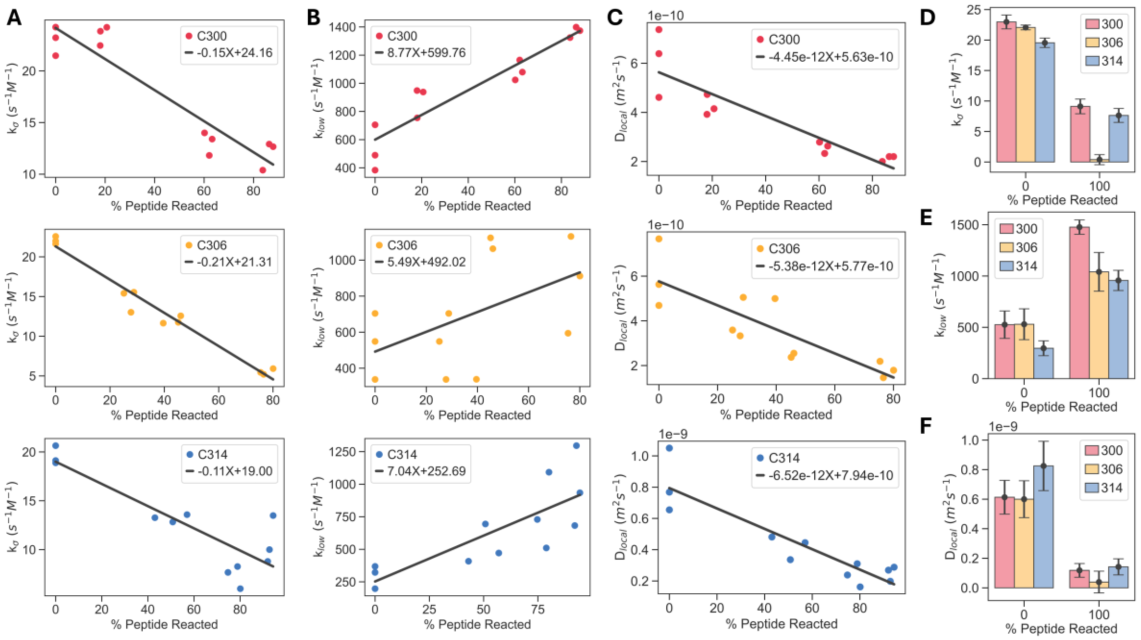


**Fig. S7.** Hydration variables over amount of reacted peptide fitted with linear line: $m * x + C$: (A) $k_\sigma$, (B) $k_{low}$, and (C) $D_{local}$. The comparison between residues using data from 0X seed and the extrapolated values at 100% peptide reacted for (D) $k_\sigma$, (E) $k_{low}$, and (F) $D_{local}$. The error bars in (D), (E), and (F) are reported as $\sqrt{(x \cdot \sigma_m)^2 + \sigma_C^2 + 2x \cdot \rho \cdot \sigma_m \cdot \sigma_C}$, based on standard error of the curve fit.

**Table S2.** Summary statistics for fibril-elongation measurements (Seed, Site 300, Site 306, Site 314). Values represent the median fibril length mean fibril length, standard deviation (SD), and sample size (n) for each condition. The data show that seed and Site 300 seeding have comparable distributions on fibril length, whereas Site 306 and Site 314 exhibit substantially higher mean fibril length.

| | Seed | 300 Seeding | 306 Seeding | 314 Seeding |
|---|---|---|---|---|
| Median (nm) | 145 | 168 | 592 | 442 |
| Mean (nm) | 183 | 166 | 593 | 449 |
| SD (nm) | 111 | 41 | 236 | 156 |
| n | 33 | 33 | 21 | 21 |

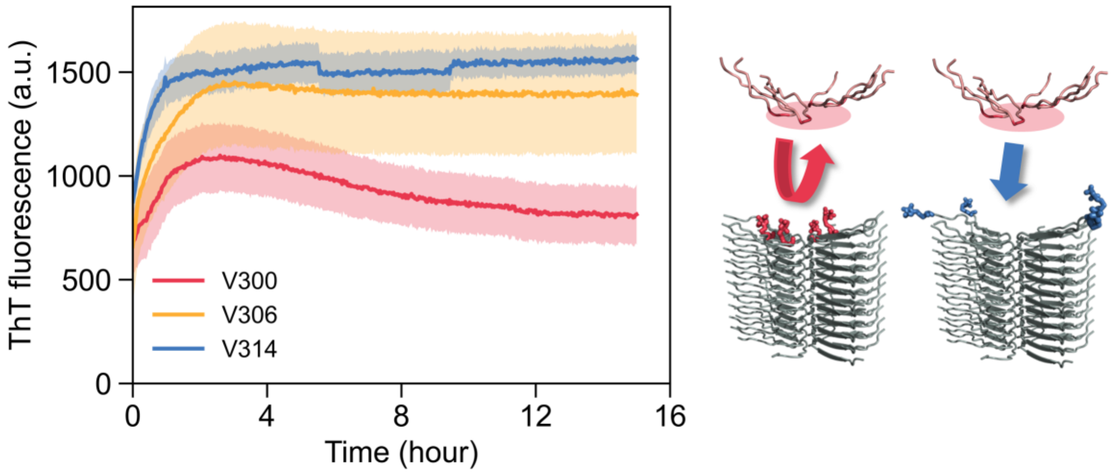


**Fig. S8.** Seeds pre-capped with spin-labeled tau remain active for docking; only capping site 300 impedes locking. Fibril seeds were pre-incubated with spin-labeled jR2R3-P301L (site 300, 306, or 314), the reaction was stopped, and the seeds were washed to remove peptide not firmly incorporated, leaving spin-labeled tau capping the fibril ends. The washed seeds were then reacted with fresh, unlabeled tau peptide and followed by ThT (site 300, orange; 306, yellow; 314, blue). All three give a similar rapid initial rise, showing the capped seeds remain competent for the initial docking of unlabeled tau peptides. Seeds capped at 306 and 314 continue to grow, indicating unlabeled tau peptide propagates past these spin-label defects, whereas seeds capped at site 300 show clearly impeded locking and net growth. Because the elongating tau peptide carries no spin label, the deficit cannot arise from the label impairing the soluble peptide and localizes the effect to occlusion of the site-300 hotspot on the fibril. This panel is interpreted qualitatively.

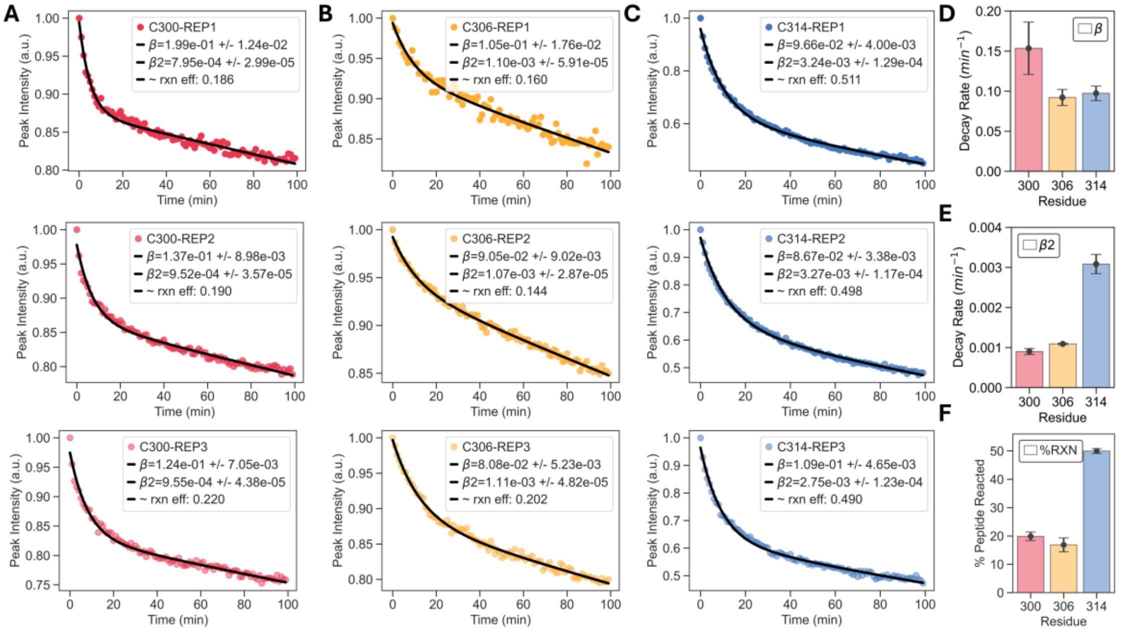


**Fig. S9.** cw-EPR central peak height over time fitted with bi-exponential decay for jR2R3-P301L 100% labeled at site (A) C300, (B) C306, and (C) C314. (D) $\beta$ comparison between residues. (E) $\beta 2$ comparison between residues. (F) Relative reaction efficiency at $T = 100\, min$ comparison between residues. The error bars in (D), (E), and (F) are reported as 1 standard deviation of the triplicates.

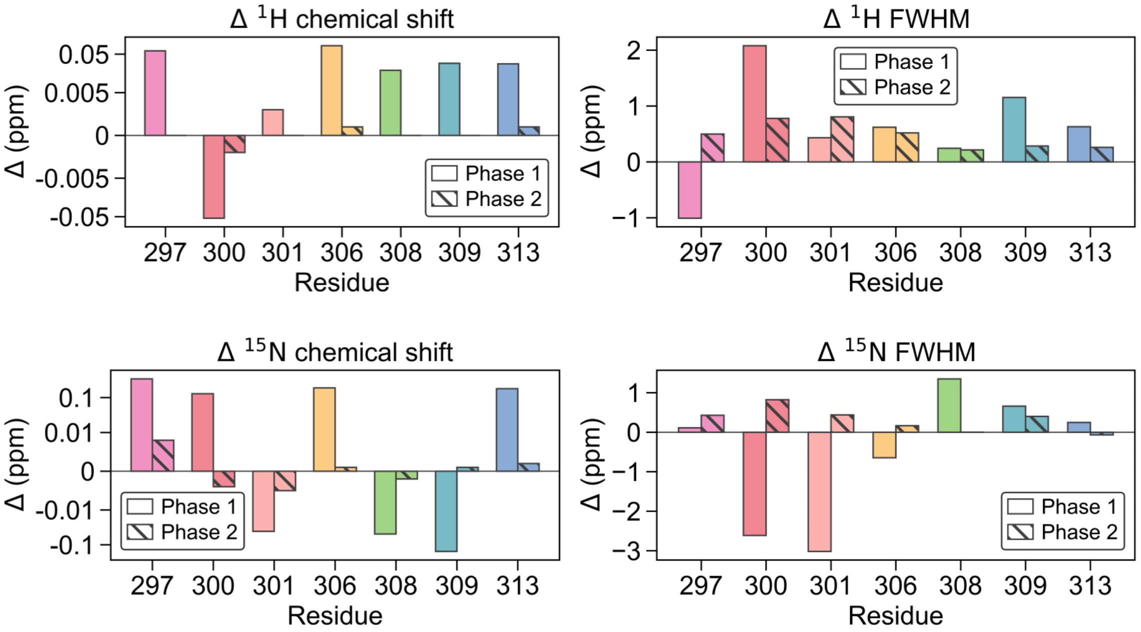


**Fig. S10. Changes in chemical shift and FWHM of the $^1$H and $^{15}$N resonances for each labeled residue across two time intervals.** Phase 1 represents the change from 0 to 1 hr, and Phase 2 represents the change from 1 to 10 hr;. Chemical shift changes are displayed on a symmetric logarithmic (symlog) y-axis to allow both phases to be visualized despite their large difference in magnitude; the axis is linear within a region of ±0.005 ppm for $^1$H and ±0.01 ppm for $^{15}$N around zero, and logarithmic beyond. FWHM changes are displayed on a linear y-axis.

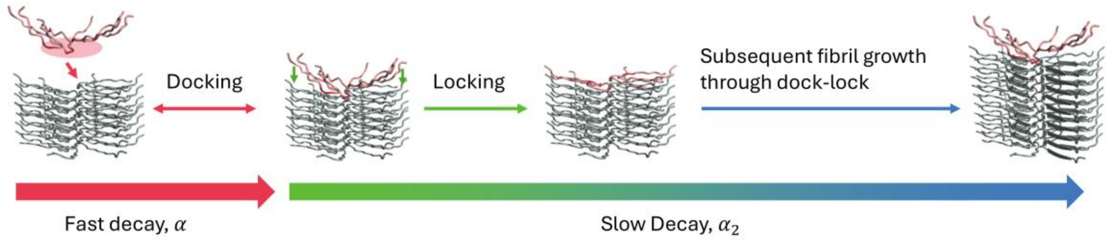


**Fig. S11. Schematic of the proposed docking-then-locking sequence at the dominant hotspot.** Soluble tau first docks site-specifically at the fibril end (the fast, residue-specific $\alpha$ phase, where the chemical-shift and linewidth changes occur), followed by locking that incorporates it into the cross-$\beta$ core and extends the fibril. Reversible, non-specific docking may occur in parallel (background) but is not required to account for the site-specific kinetics; the docking→locking sequence is the interpretation most consistent with the data and is not presented as the only possible mechanism.

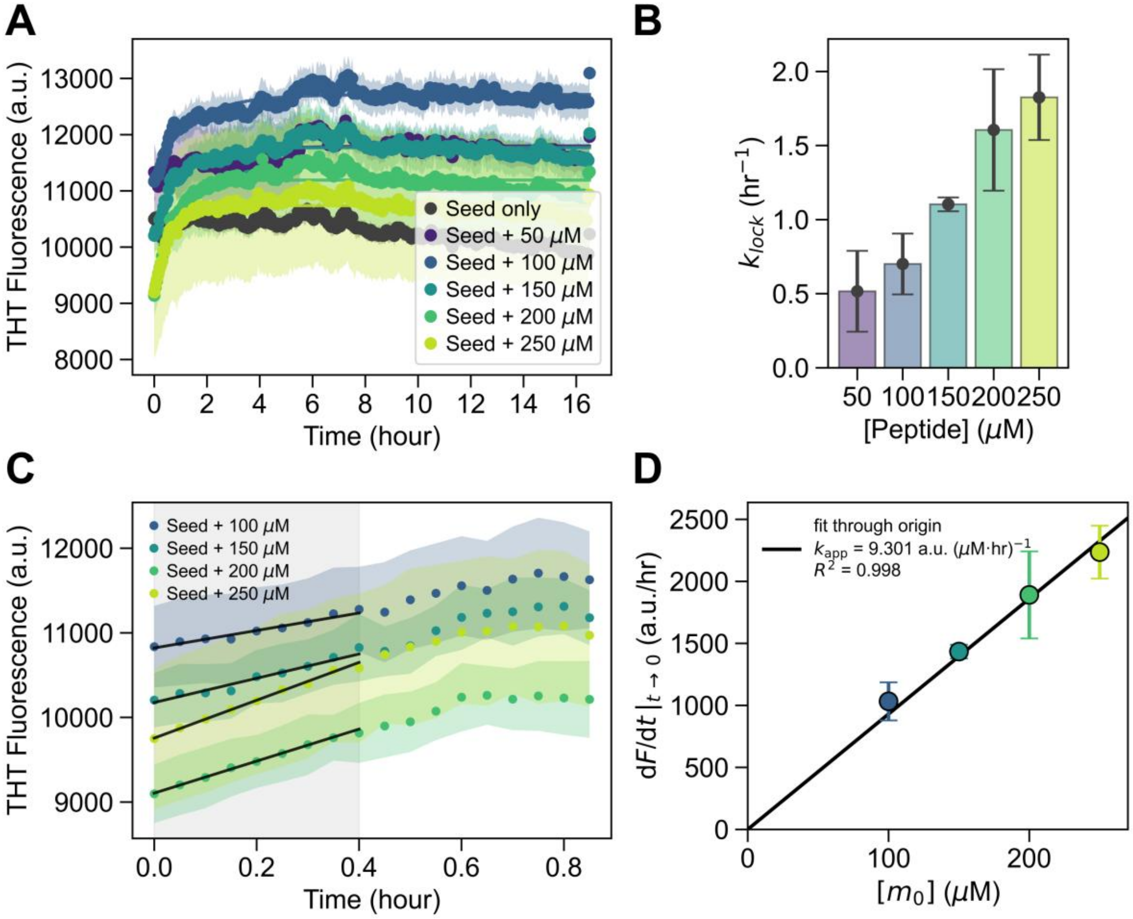


**Fig. S12. Peptide-concentration dependence of ThT elongation kinetics (50–250 μM peptide, 100 μM seed), analyzed two ways.** (A) ThT traces at each peptide concentration (mean ± 1 SD, n = 3). (B) Single-exponential, lock-limited analysis: each trace was fit to $F(t) = F_0 + F\_max(1 - exp(-k\_lock \cdot t))$, the $k\_dock \gg k\_lock$ limit of the dock-and-lock scheme (Esler et al., 2000) (24), and the fitted k_lock is plotted against peptide concentration; k_lock rises with concentration. Under sub-saturating conditions (peptide below K_D), k_lock is an apparent rate carrying the peptide dependence of docked-complex occupancy rather than the intrinsic locking constant. (C) Initial-rate analysis: the first 0.4 hr of the traces (100–250 μM; shaded) overlaid with linear initial-slope fits (black), from which dF/dt|_(t→0) was obtained. (D) Per-condition mean initial slopes (n = 3) plotted against initial peptide concentration $m_0$, with a through-origin fit giving the apparent elongation rate constant $k\_app = 2k_+P_0 = 9.30$ a.u.·$\mu M^{-1} \cdot hr^{-1}$ ($R^2 = 0.998$); 50 μM was excluded because its initial-rate signal fell within baseline drift. Both analyses show the apparent elongation rate increasing with peptide concentration, as expected when the elongation-competent pool scales with monomer concentration; absolute $k_+$ was not extracted because the fibril-end concentration $P_0$ in the sonicated seed stock was not independently determined.

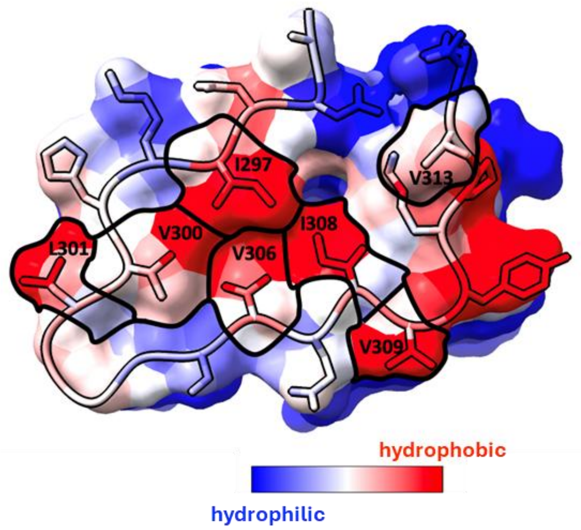


**Fig. S13.** INDUS hydrophobicity landscape of every residue on jR2R3-P301L.
